# The yeast peroxisomal proteome at absolute quantitative scale

**DOI:** 10.1101/2025.10.20.683451

**Authors:** Hirak Das, Silke Oeljeklaus, Renate Maier, Julian Bender, Bettina Warscheid

## Abstract

Peroxisomes are dynamic organelles vital for lipid metabolism and redox homeostasis. In *Saccharomyces cerevisiae*, the expression of peroxisomal proteins is tightly regulated in response to metabolic conditions. Here, we provide the first absolute quantification of the yeast peroxisomal proteome under peroxisome-inducing (oleate) and fermentative (glucose) conditions using a label-free mass spectrometry approach. We determined protein copy numbers for ∼4,500 proteins, including 99 peroxisomal and peroxisome-associated proteins. Our data reveal that the peroxisomal proteome is ∼3-fold higher abundant in oleate-grown cells, constituting 2.8% of the total proteome (corresponding to 2.01 x 10^6^ protein copies) compared to 0.8% (6.67 x 10^5^ protein copies) in glucose, reflecting the necessity for peroxisomal functions such as fatty acid beta-oxidation. Enzymes of the beta-oxidation and glyoxylate cycle showed up to >500-fold higher abundance in oleate. In contrast, core components of the peroxisomal protein import machinery (e.g., Pex5, Pex14) exhibited only moderate changes (∼2- to 8-fold). In addition to metabolic enzymes and components of the peroxisomal protein import pathways, we provide copy number data for proteins involved in cellular stress response, peroxisome proliferation, division and organization, peroxisome-associated membrane contact sites, and metabolite transporter. Taken together, our dataset offers a quantitative framework of peroxisomal remodeling under different metabolic conditions and highlights the organelle’s adaptive flexibility, providing a valuable resource for future studies on peroxisome biology.

## Introduction

Peroxisomes are ubiquitous single membrane-bound organelles present in nearly all eukaryotic cells. They are key players in cellular metabolism, in particular in lipid metabolism (Waterham et al. 2016) and H_2_O_2_ metabolism (Lismont et al. 2019), but they have also been implicated in non-metabolic functions such as redox homeostasis (Ferreira et al., the immune response to viral infections, inflammation, cancer and age-dependent diseases (reviewed in Zalckvar and Schuldiner 2022). Furthermore, peroxisomes are highly dynamic and adjust their number, size, and protein repertoire – the peroxisomal proteome – to the metabolic, environmental, or developmental requirements of the cell. In the yeast *Saccharomyces cerevisiae*, for example, peroxisome growth and proliferation are increased when fatty acids (FAs) such as oleic acid are used as carbon source (Veenhuis et al. 1987; Gurvitz and Rottensteiner 2006). Under these metabolic conditions, the transcription of genes required for FA degradation via the beta-oxidation pathway, which in *S. cerevisiae* is exclusively compartmentalized in peroxisomes, is activated (Hiltunen et al. 2003). In contrast, when cells are grown under fermentative conditions on glucose, the expression of genes required for peroxisomal functions is repressed (Gurvitz and Rottensteiner 2006). Accordingly, glucose-grown cells typically harbor only 1 - 2 peroxisomes per cell, while oleate-grown cells have higher, varying numbers of larger peroxisomes (up to ∼ 20 per cell with an average of ∼ five) (Veenhuis et al. 1987; Erdmann and Blobel 1995; Kuravi et al. 2006). *S. cerevisiae* has been used for decades as prime model organism in peroxisome research, not only to identify the majority of peroxisomal biogenesis factors (i.e., peroxins or Pex for short), but also to decipher fundamental mechanisms of peroxisome biology including the import of peroxisomal membrane and matrix proteins (Rudowitz and Erdmann 2023), peroxisome proliferation and division (Schrader et al. 2016), inheritance (Knoblach and Rachubinski 2016), or the formation of membrane contact sites with other organelles (Silva et al. 2020). For an advanced understanding of peroxisomal processes and functions in a cellular context, knowledge of the absolute abundance of the individual proteins that mediate the processes provides an additional, in-depth level of information, as it was demonstrated for the mitochondrial proteome of yeast and human cells (Morgenstern et al. 2017, 2021; Pfanner et al. 2019).

Several studies have reported protein copy number data for *S. cerevisiae* grown under different conditions (summarized by Ho et al. 2018). However, none of the absolute quantitative studies to date included peroxisome-inducing growth conditions. To obtain a detailed quantitative understanding of the yeast peroxisomal proteome, we estimated protein copy numbers for yeast cells grown under peroxisome-inducing and fermentative conditions using oleic acid and glucose as carbon source. For this, we followed a label-free quantitative mass spectrometry (MS)-based approach and applied the “proteomic ruler” method (Wiśniewski et al. 2014). We report copy numbers for ∼4,500 proteins including 99 proteins associated with different aspects of peroxisome biology. Our absolute quantitative data reveal that peroxisomal proteins constitute only 2.8% and 0.8% of the overall protein copy numbers in oleate- and glucose-grown cells, respectively. Under peroxisome-inducing condition, enzymes of the fatty acid beta-oxidation and the glyoxylate cycle drastically increase up to >500 fold, while the increase in proteins of the peroxisomal import machineries is moderate with up to 8-fold. We also present copy number data for proteins associated with other important peroxisomal processes including cellular stress response, proliferation, division, and organization, organellar membrane contact sites and metabolite transport. Overall, our depiction of the peroxisomal proteome at an absolute quantitative scale provides interesting insights into peroxisome biogenesis and its diverse functions in a metabolic and cellular context.

## Materials and Methods

### Cultivation of yeast cells

Experiments were performed using the *S. cerevisiae* strain BY4741 (MATa his3Δ1 leu2Δ0 met15Δ0 ura3Δ0; Brachmann et al. 1998). Cells were grown overnight at 30°C and 160 rpm in synthetic complete (SC) medium containing 0.17% (w/v) yeast nitrogen base (YNB), 0.5% (w/v) ammonium sulfate, 0.3% (w/v) glucose, and selected amino acids and nucleobases (pH 6.0) (Schummer et al. 2017). Fresh SC medium was inoculated with an aliquot of the overnight culture at an OD_600_ of 0.2, and cells were cultivated for further 8 h as described above. For growth under peroxisome-proliferating conditions, cells were shifted to YNO medium (0.17% [w/v] YNB, 0.5% [w/v] ammonium sulfate, 0.1% [v/v] oleic acid, 0.05% [v/v] Tween 40, and selected amino acids and nucleobases; pH 6.0) (Schummer et al. 2017). For growth under fermentative conditions, SC medium was supplemented with 2% (w/v) glucose. Cells were cultured at 30°C and 160 rpm for further 16 h, harvested by centrifugation (10 min at 7,000 x g and room temperature), and washed twice with deionized water.

### Preparation of whole cell lysates and proteolytic in-solution digestion

Yeast cells (400 µg per replicate) were resuspended in 500 µl of lysis buffer (8 M urea, 75 mM NaCl, 50 mM Tris/HCl, 1 mM EDTA, pH 8.0). 300 mg of glass beads (425-600 µm, Sigma-Aldrich/Merck) were added, and the cells were mechanically disrupted by two cycles of bead beating for 4 min at 4,000 rpm using a MiniLys homogenizer (Bertin Technologies, France) with at least 4 min cooling on ice between the cycles. Glass beads, cell debris and unbroken cells were removed by centrifugation (5 min at 15,000 x g and 4°C), and the protein concentration of the lysates was adjusted to 1 µg/µl using digestion buffer (8 M urea in 50 mM ammonium bicarbonate). Cysteine residues were reduced using Tris(2-carboxyethyl)phosphine at a final concentration of 5 mM and incubation for 30 min at 37°C, followed by alkylation of free thiol groups with 2-chloroacetamide (50 mM final concentration; 30 min at room temperature). The reaction was quenched by adding DTT (25 mM final concentration). Samples were diluted with 50 mM ammonium bicarbonate to reach a final urea concentration of 1.6 M. Trypsin was added at a protease-to-protein ratio of 1:50 and proteins were digested overnight at 37°C with slight agitation. Proteolysis was stopped by addition of trifluoroacetic acid (TFA; 1% final concentration). Peptide mixtures were desalted using C18 cartridges (3M Empore, St. Paul, USA) according to the manufacturer’s protocol and dried in vacuo.

### High-pH reversed-phase peptide fractionation

Tryptic peptides corresponding to 300 µg of protein were fractionated by high-pH reversed-phase chromatography (Delmotte et al. 2007) using an NX 3u Gemini C18 column (150 mm x 2 mm, particle size 3 µM, pore size 110 Å; Phenomenex, Aschaffenburg, Germany) and a binary solvent system consisting of 10 mM ammonium hydroxide, pH 10, (solvent A) and 90% (v/v) acetonitrile (ACN)/10 mM ammonium hydroxide (solvent B). Dried peptides were dissolved in 400 µl of solvent A by sonication in an ultrasonic bath for 5 min. Insoluble material was removed by centrifugation (5 min at 12,000 x g), and supernatants were filtered using a 0.2 µm PTFE membrane syringe filter (Phenomenex) and loaded onto the column using an Ultimate^TM^ 3000 HPLC system (Thermo Fisher Scientific, Dreieich, Germany) operated at a flow rate of 200 µl/min and a column temperature of 40°C. Peptides were loaded for 2 min at 1% solvent B and separated using a gradient ranging from 1 – 50% B in 20 min, followed by 50 – 78% B in 1 min and 1 min at 78% B. The column was re-equilibrated with 100% A for 6 min. Fractions were collected in 50-s intervals from minute 1 to minute 28 in a concatenated manner, resulting in a total of eight fractions. Peptides were dried in vacuo and stored at -80°C until used for liquid chromatography-mass spectrometry (LC-MS) analysis.

### Mass spectrometry

Dried peptides were dissolved in 33 µl of 0.1% (v/v) TFA by sonication for 3 min. Insoluble material was removed by centrifugation (6 min at 12,000 x g), supernatants were transferred into fresh glass vials, and 5 µl of each fraction were used for LC-MS analysis using an UltiMate^TM^ 3000 rapid separation liquid chromatography system (RSLCnano, Thermo Fisher Scientific, Dreieich, Germany) coupled to a Q-Exactive^TM^ Plus mass spectrometer (Thermo Fisher Scientific, Bremen, Germany). Peptides were washed and preconcentrated on µPAC^TM^ C18 trapping columns (10 mm x 2 mm inner diameter; PharmaFluidics, Ghent, Belgium) at a flow rate of 10 µl/min and separated on a µPAC^TM^ C18 analytic column (500 mm x 0.3 mm; PharmaFluidics) at a flow rate of 0.3 µl/min and 40°C. Peptides were separated using a solvent system consisting of 0.1% (v/v) formic acid (solvent A) and 86% (v/v)/0.1% (v/v) formic acid (solvent B). Peptides were loaded onto the trap columns for 3 min at 1% solvent B and eluted using the following gradient: 1 – 20% B in 103 min, 20 – 42% B in 50 min, 42 – 95% B in 2 min and 5 min at 95% B. High resolution full MS scans were acquired in a mass-to-charge (*m/z*) range of 375 to 1,700 and at a resolution of 70,000 (at *m/z* 200). The automatic gain control was set to 3 x 10^6^ ions at a maximum ion injection time of 60 ms. For fragmentation of the 12 most intense peptide ions (z ≥ 2) by high energy collision dissociation, the normalized collision energy was set to 28%. Fragment ions were recorded at a resolution of 35,000 with an automatic gain control of 1 x 10^5^, a maximum injection time of 120 ms, and a dynamic exclusion time of 45 s.

### Data analysis

Proteins were identified using MaxQuant/Andromeda (version 2.6.6.0; (Tyanova et al. 2016a). MS/MS data were searched against all entries in the “orf_trans_all” fasta file downloaded from the *Saccharomyces* Genome Database (SGD; https://www.yeastgenome.org/, version from April 2024). The database search was performed with trypsin/P as proteolytic enzyme, N-terminal acetylation and methionine oxidation as variable modifications, carbamidomethylation of cysteine residues as fixed modification, and a maximum of three missed cleavage sites. The minimum number of peptides and unique peptides (minimum length of seven amino acids) required for protein identification were set to one each. A false discovery rate of 0.01 was applied to both peptide and protein identifications. “Match-between-runs” was enabled, allowing matching only between samples generated from cells grown under the same conditions.

### Estimation of protein copy numbers per cell and further data analysis

Copy numbers for proteins identified in oleate- and glucose-grown cells (four biological replicates each) were estimated using Perseus (version 2.1.5.0; Tyanova et al. 2016b) and the Perseus plugin “Proteomic Ruler” (version 1.6.2; https://maxquant.net/perseus_plugins/). The “proteomic ruler” method (Wiśniewski et al. 2014) can be used for organisms and cellular systems whose ploidy and genome size is known. It is based on the fact that cellular DNA is bound to histones in a fixed stoichiometry. Consequently, the sum of all MS signals from histone-derived peptides can be used as a “ruler” to determine the DNA content of the sample. Since the amount of DNA is proportional to the number of cells, the method facilitates the calculation of the total cellular protein mass. The intensities of all other proteins identified in the sample can be scaled accordingly to estimate the mass of each protein per cell and, which can be converted to numbers of proteins (i.e., copy numbers) per cell based on the molecular mass of the protein. The Perseus-based copy number estimation was carried out using MaxQuant MS intensities, the protein sequences present in the proteome reference set for *S. cerevisiae* (strain ATCC 204508/S288c; proteome ID UP000002311 containing 6,067 entries), and the following settings: proteolytic enzyme, trypsin/P; minimum/maximum peptide length, 7/30; detectability correction was enabled, with number of theoretical peptides (trypsin/P, 7 – 30) as correction factor; and ploidy, 1.

The results table from Perseus was parsed in Python using the autoprot package (Bender et al. 2024). Protein copy numbers of replicates were log_2_-transformed, and mean log_2_fold-changes between cells grown on oleate versus glucose across all replicates per growth condition as well as statistical significance of these changes were calculated using the R package limma (Smyth 2004) interfaced through autoprot. Results of the LC-MS data analysis and copy number estimation are provided in **Supplementary Table S1**. GO term enrichment analyses were performed using g:Profiler (Kolberg et al. 2023) via autoprot. Results are provided in **Supplementary Table S2**.

### Data availability

The mass spectrometry proteomics data have been deposited to the ProteomeXchange Consortium (Deutsch et al. 2023) via the PRIDE (Perez-Riverol et al. 2025) partner repository and are accessible using the dataset identifier PXD069526.

## Results and Discussion

### Estimation and assessment of cellular protein copy numbers

To quantitatively explore the peroxisomal proteome of baker’s yeast and its differences between fermentative and peroxisome-inducing growth conditions, we cultivated yeast cells in glucose- and oleate-containing medium. Tryptic peptide mixtures obtained from whole cell lysates were separated by high-pH reversed-phase fractionation into 8 fractions per sample to facilitate high coverage of the peroxisomal proteome, followed by LC-MS analysis (**Fig. 1a**). Experiments were performed in four independent replicates per growth condition. We used the “proteomic ruler” method (Wiśniewski et al. 2014) to estimate protein copy numbers based on MS intensities (for details, see Methods). As a result, we determined absolute protein copy numbers for 4,392 proteins for oleate- and 4,388 proteins for glucose-grown cells (**Table S1**). 4,332 proteins were identified under both metabolic conditions, while 60 proteins were only identified in oleate- and 56 proteins only in glucose-grown cells (**Fig. S1a**; **Table S1**). Pearson correlation coefficients of 0.96 - 0.98, determined for log_2_-transformed protein copy numbers, reveal a high reproducibility across all replicates (**Fig. S1b**).

**Fig. 1.**
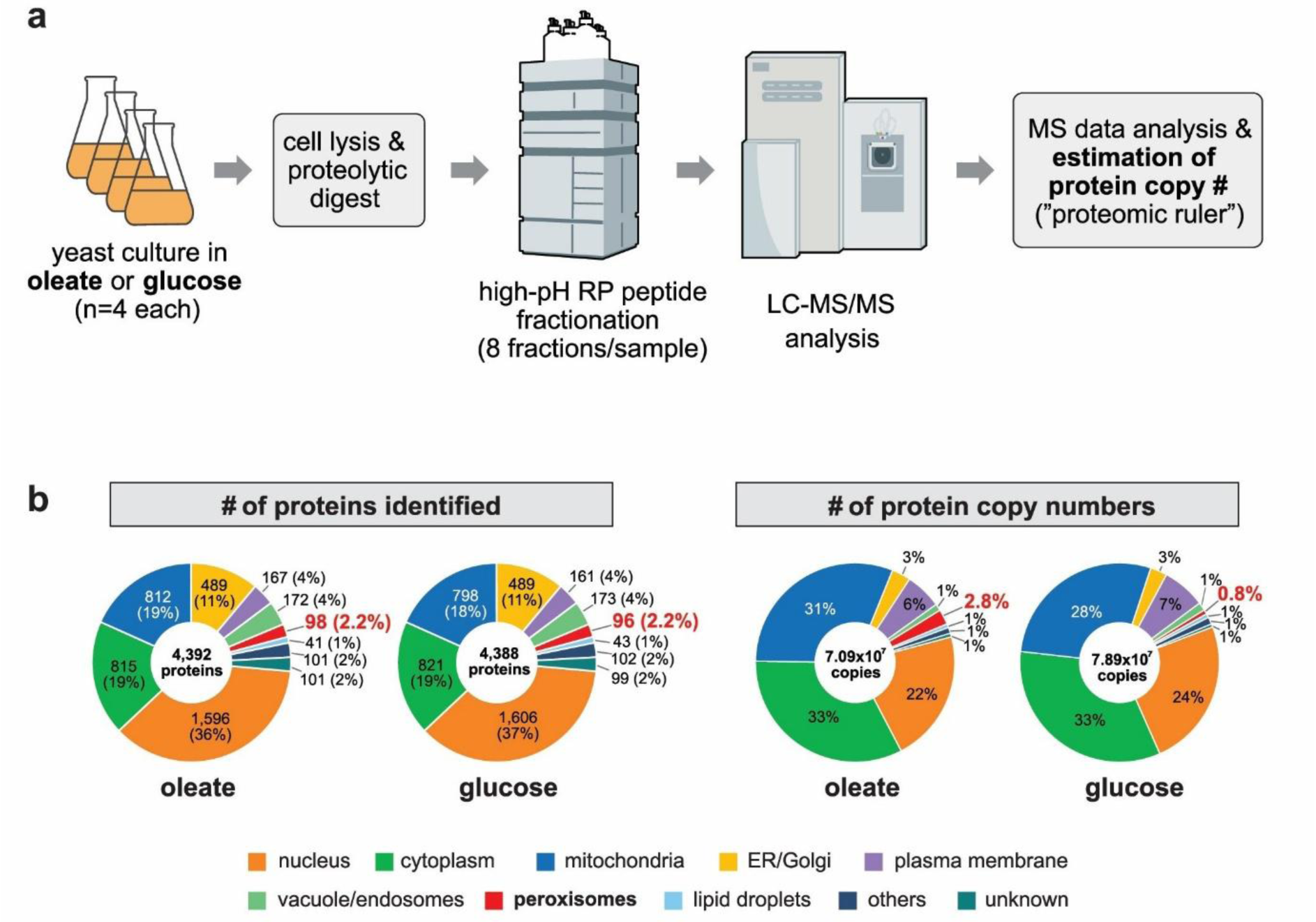
Quantification and analysis of cellular protein copy numbers. **a.** Workflow used for the mass spectrometry-based estimation of absolute protein copy numbers in *S. cerevisiae* cells grown under peroxisome-proliferating (oleate) or fermentative (glucose) conditions. RP, reversed-phase; LC-MS/MS, liquid chromatography-tandem mass spectrometry. **b.** Carbon source-dependent abundance of cellular sub-proteomes based on the number of proteins identified (left) and protein copy number (right). Non-peroxisomal proteins were assigned to the respective subcellular localization based on Gene Ontology annotations. The peroxisomal proteome was defined based on literature (van Roermund et al. 2021; Yifrach et al. 2022; Kosir et al. 2025; Chen et al. 2025).

The proteome of *S. cerevisiae* is currently estimated to comprise 6,052 proteins (*Saccharomyces* Genome Database, http://sgd.yeastgenome.org; dubious ORFs excluded; see version 2024-06-13 of the orf_trans.fasta.gz file), of which our combined copy number dataset from glucose- and oleate-grown cells covers 4,448 proteins (73.5%) (**Table S1**). The distribution of identified proteins to different subcellular locations shows virtually no difference between both metabolic conditions: 85% of all identified proteins were assigned to the nucleus, representing the largest fraction (36% for oleate-, 37% for glucose-grown cells), the cytosol/cytoplasm (19% each), mitochondria (19% and 18%, respectively), and the ER/Golgi (11% each) (**Fig. 1b, left panel**). The remaining 15% are proteins of the plasma membrane, vacuole/endosomes, peroxisomes, lipid droplets, proteins whose localization is not further specified here (“others”) and whose localization is currently still unknown. We identified 99 (83.2%) of the presently reported 119 peroxisomal and peroxisome-associated proteins (van Roermund et al. 2021; Yifrach et al. 2022; Kosir et al. 2025; Chen et al. 2025) (**Table 1**), for simplicity’s sake in the following collectively referred to as peroxisomal proteins when a distinction is not relevant. Irrespective of the growth condition, peroxisomal proteins equally accounted for 2.2% of all identified proteins, i.e., 98 proteins in oleate- and 96 proteins in glucose-grown cells (**Fig. 1b, left**). 94 peroxisomal proteins were identified under both metabolic conditions, three proteins (Pex12, Pex15, Pex9) were only identified in oleate-, and one (Gto1) only in glucose-grown cells (**Table 1**), indicating that the yeast peroxisomal proteome is expressed despite different metabolic requirements of the cells.

**Table 1.**
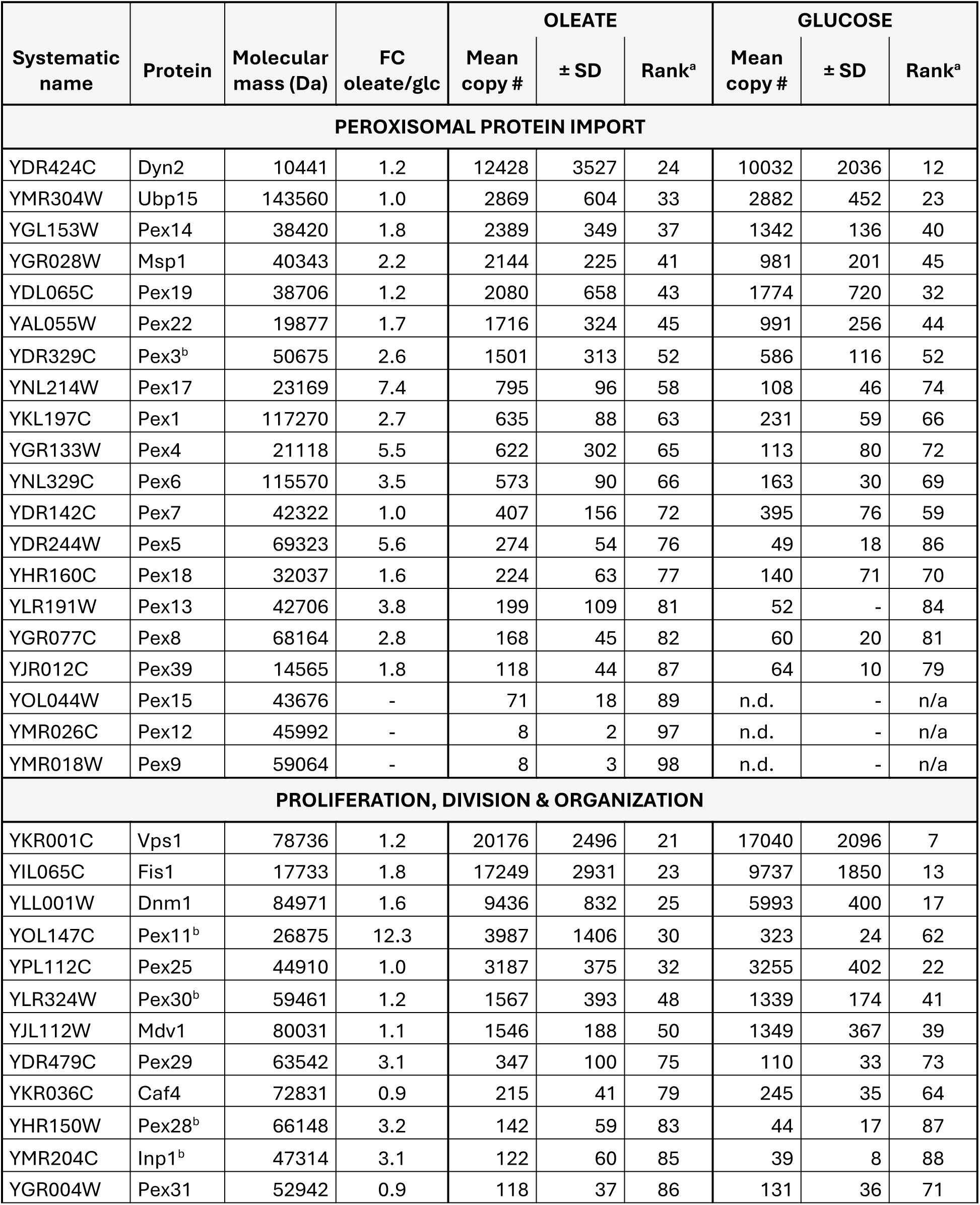
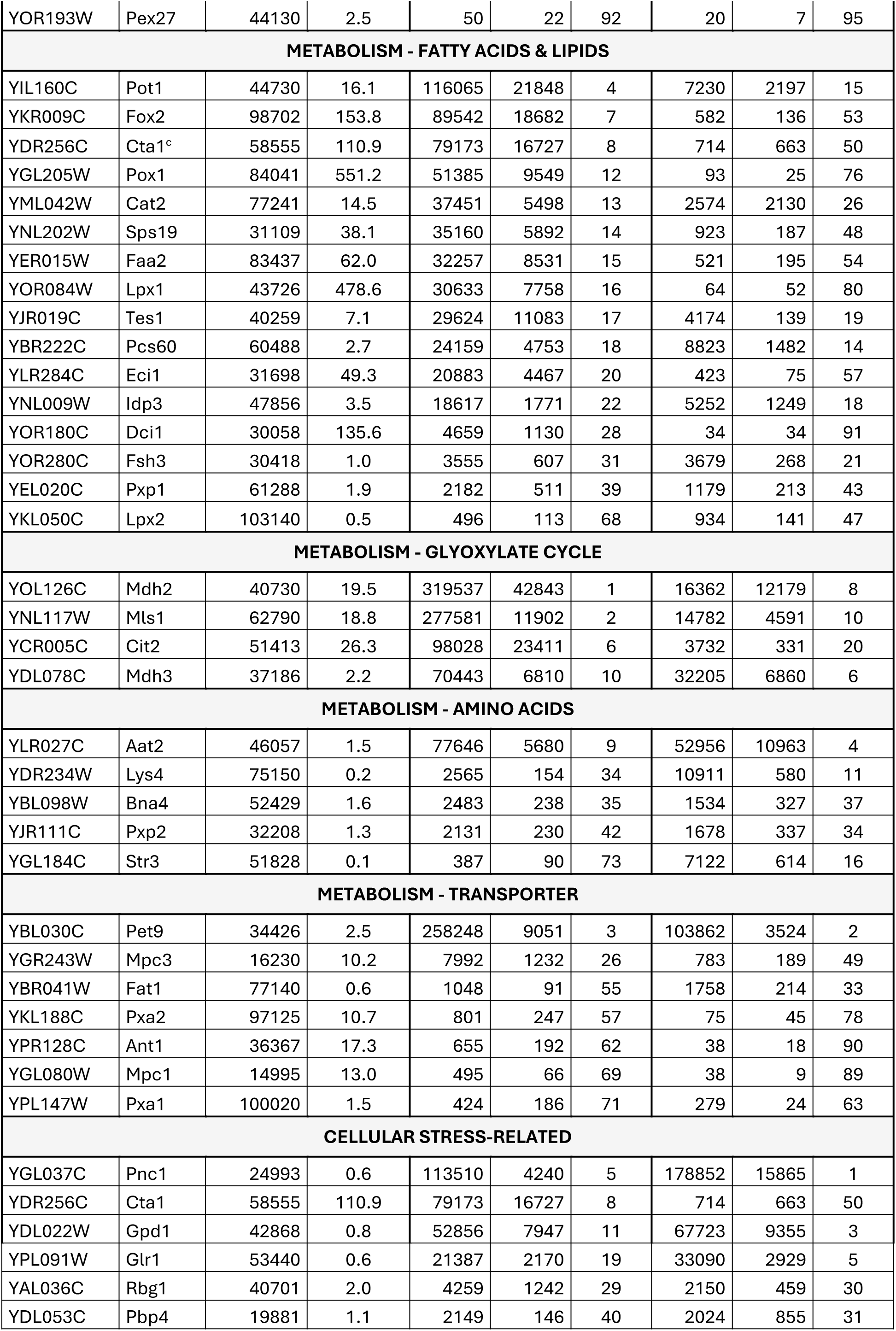
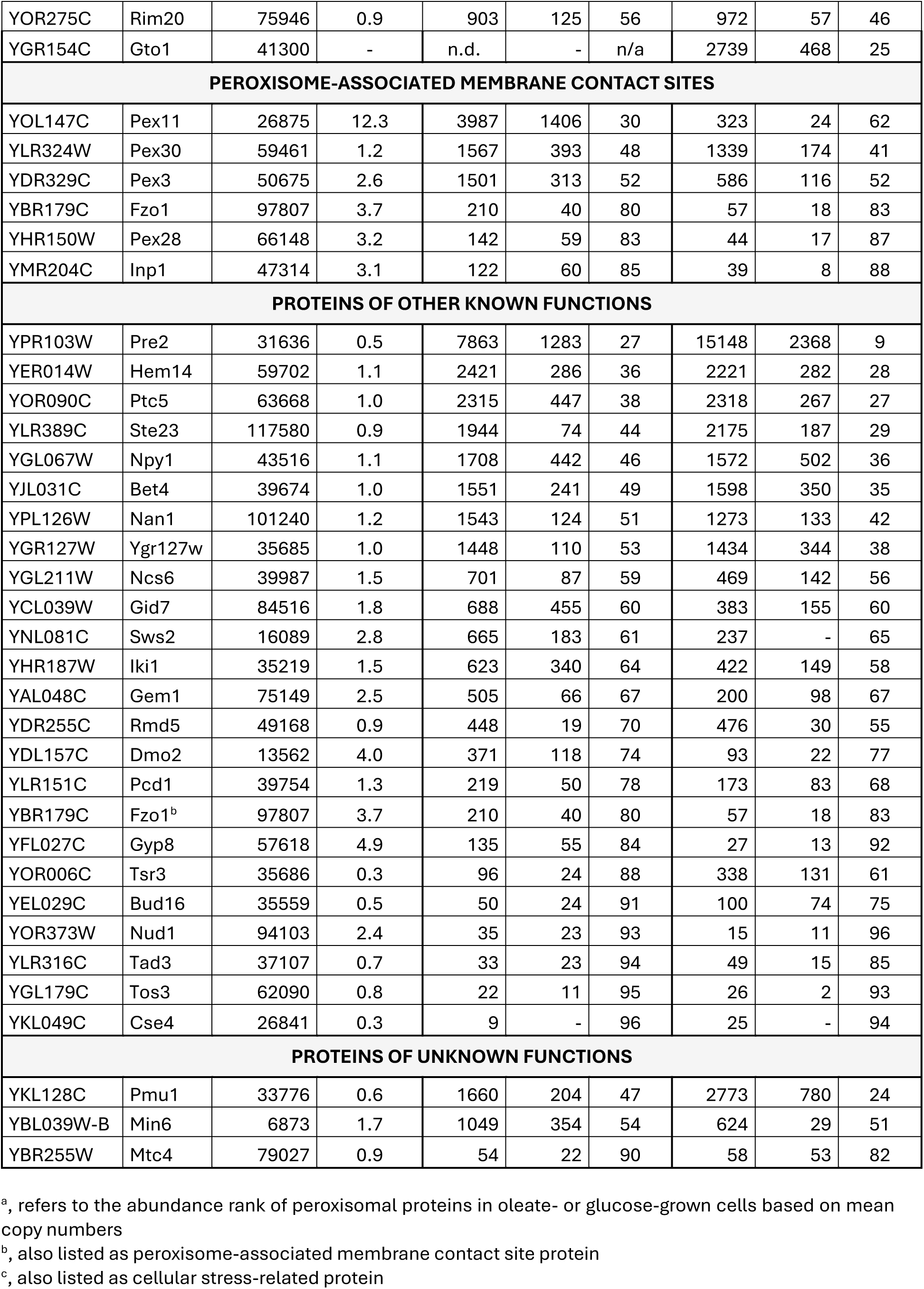
Relative and absolute quantification of peroxisomal proteins identified in yeast cells grown under peroxisome-proliferating (oleate) and fermentative conditions (glucose). FC, fold-change; glc, glucose; SD, standard deviation; n.d., not determined; n/a, not applicable.

In terms of absolute protein abundance, overall cellular protein copy numbers were in a similar dimension for oleate- and glucose-grown cells: 7.09 x 10^7^ (oleate) *versus* 7.89 x 10^7^ (glucose) protein copies (**Fig. 1b, right panel**). Cytosolic/cytoplasmic, mitochondrial and nuclear proteins accounted for 86% and 85% of the total cellular proteome in oleate- and glucose-grown cells, but with different fractions compared to protein identification numbers. Of all the main subcellular niches, only peroxisomes showed a considerable difference in protein copy numbers: in cells grown on oleate, the total copy number for peroxisomal proteins was ∼3-fold higher than in glucose-grown cells, increasing from 0.8% on glucose to 2.8% of the total cellular proteome on oleate (**Fig. 1b, right**). This finding is in accordance with a higher demand for peroxisomal functions, in particular fatty acid beta-oxidation, when cells are supplied with oleic acid as sole carbon source. Our data show that peroxisomal proteins account for only a small fraction of the total yeast proteome, even under peroxisome-inducing conditions. Moreover, many proteins associated with peroxisome functions generally have dual or multiple subcellular localizations, with often only a small part present in or at peroxisomes, depending on the metabolic or environmental condition of the cell. Of note, the experimental design of our study does not allow for monitoring the subcellular localization of proteins under the different growth conditions and, thus, does not provide information about the fraction of multi-localized proteins in peroxisomes. Nevertheless, we can conclude that despite the importance of peroxisomes in cellular metabolism, their proteome represents just a tiny fraction of the total cellular proteome in terms of protein number and absolute abundance.

### Growth-dependent differences in the relative abundance of cellular proteins

To reveal differences in protein abundance between oleate- and glucose-grown cells at the single protein level, we plotted mean log_2_ oleate-*versus*-glucose copy number ratios for all 4,332 proteins identified under both conditions against the corresponding -log_10_-transformed adjusted p-value (**Fig. 2a**). A total of 505 proteins were significantly higher abundant in oleate-grown cells (p-value < 0.05; oleate/glucose ratio ≥ 1.5), while 625 proteins had higher levels under glucose condition (p-value < 0.05; ratio ≤ 0.6667). Gene Ontology (GO) term enrichment analysis revealed that peroxisomal (matrix and membrane) and mitochondrial (intermembrane space, inner membrane and matrix) proteins are overrepresented in the pool of proteins with higher abundance in oleate, while cytosolic proteins are overrepresented in glucose condition (**Fig. 2b, Table S2**; GO domain “Cellular Component”, GOCC). This is also reflected by the distribution of mitochondrial, peroxisomal and cytosolic/cytoplasmic proteins in the respective volcano plots (**Fig. 2c**). GOCC terms related to nucleus, ER/Golgi, plasma membrane, vacuole/endosomes, and lipid droplets were not enriched, although individual proteins do show significant differences between oleate- and glucose-grown cells (**Fig. S2a**).

**Fig. 2.**
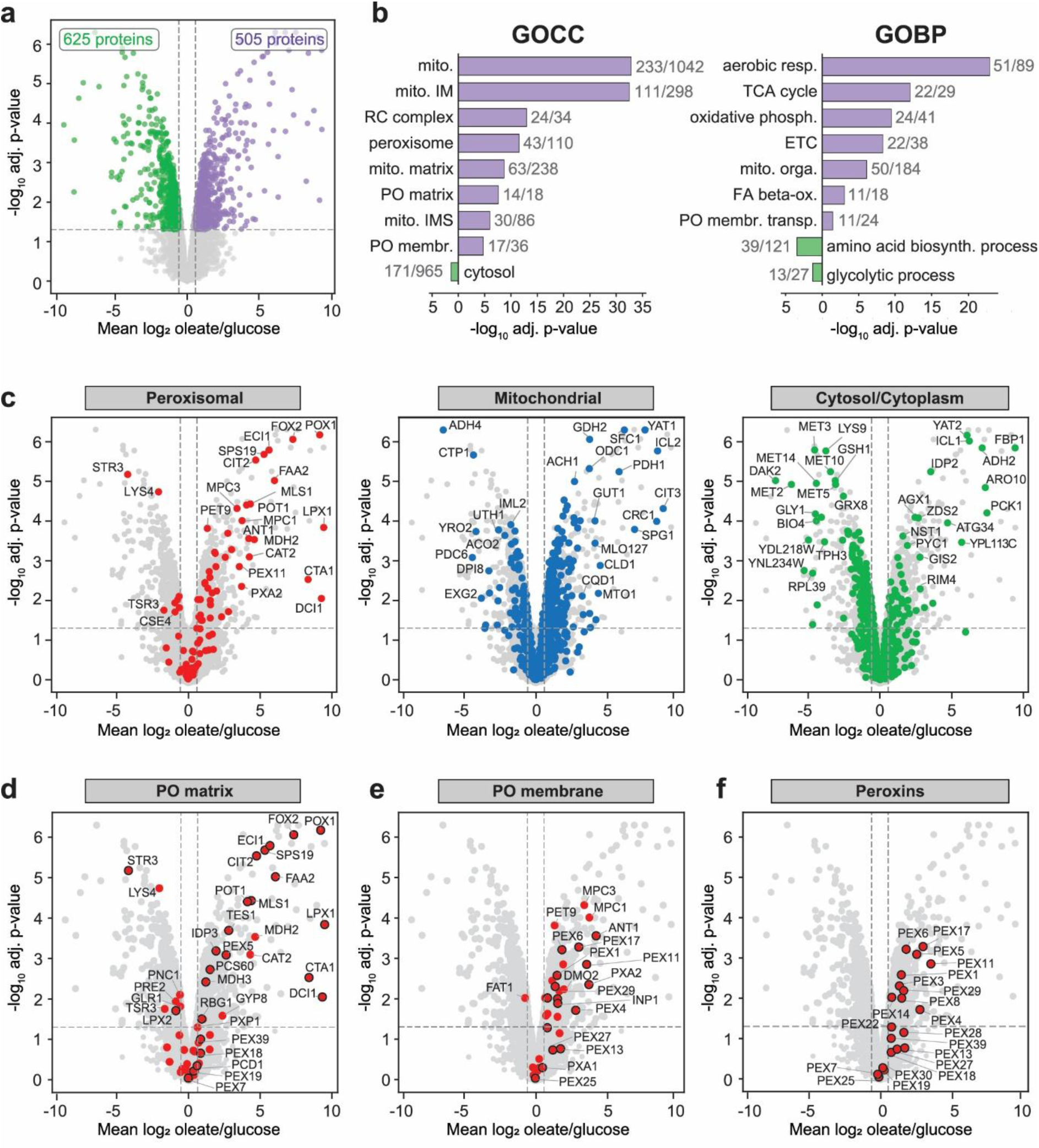
Differences in protein abundance in oleate- versus glucose-grown cells. **a.** Global differences in protein copy numbers determined for individual proteins in oleate- and glucose-grown cells. Mean log_2_ oleate/glucose copy number ratios were plotted against the corresponding -log_10_-transformed Benjamini-Hochberg-adjusted (adj.) p-value. Dashed horizontal and vertical lines indicate a p-value of 0.05 and oleate/glucose ratios of 0.6667 and 1.5, respectively. **b.** Gene Ontology (GO) term overrepresentation analysis of proteins with an oleate/glucose ratio ≥ 1.5 or ≤ 0.6667 and a p-value of < 0.05 for the domains “Cellular Component” (GOCC) and “Biological Process” (GOBP). Shown are selected terms with an adjusted p-value of < 0.05. Numbers next to bars indicate number of proteins assigned to the term in the dataset tested and number of proteins with this term in the entire dataset. biosynth., biosynthesis; ETC, electron transport chain; FA, fatty acid; IM, inner membrane; IMS, intermembrane space; membr., membrane; mito., mitochondrial; orga., organization; ox., oxidation; phosph., phosphorylation; PO, peroxisome; RC, respiratory chain; resp., respiration; TCA, tricarboxylic cycle; transp., transport. **c – f.** Same plot as in (**a**) highlighting proteins of the indicated subcellular localization/category. Proteins exclusively or predominantly localized in peroxisomes are indicated with a black border (**d** – **f**).

For *S. cerevisiae*, glucose is the preferred source of energy and building blocks for cell proliferation, although it can grow on a wide range of carbon sources. When glucose is available, the yeast preferentially metabolizes glucose via glycolysis and fermentation instead of mitochondrial respiration, even if oxygen is available. Utilization of alternative carbon sources, including fatty acids, and mitochondrial processes are repressed, which is often, but not exclusively, regulated at transcriptional level (Kayikci and Nielsen 2015). Accordingly, proteins assigned to biological processes such as aerobic respiration, tricarboxylic acid cycle, oxidative phosphorylation or the peroxisomal fatty acid beta-oxidation show significantly higher levels in oleate-grown cells, whereas proteins involved in glycolysis and amino acid biosynthesis are more abundant in glucose-grown cells (**Fig. 2b, Table S2**; GO domain “Biological Process”, GOBP). Selected examples for non-peroxisomal genes/proteins regulated by glucose-repression are the mitochondrial protein glycerol kinase Gut1, which converts glycerol to glycerol-3-phosphate (Grauslund et al. 1999), the cytosolic phosphoenolpyruvate carboxykinase Pck1, a key enzyme of the gluconeogenesis (Proft et al. 1995), and the monocarboxylate/proton symporter Jen1, which is located in the plasma membrane and mediates the import of different carbon sources such as lactate, pyruvate, and acetate (Chambers et al. 2004). They all show high oleate-to-glucose abundance ratios (**Figs. 2c, S2a; Table S1**).

### Remodeling of the peroxisomal proteome under oleate

The majority of peroxisomal proteins with significant differences in protein levels were increased in oleate-grown cells (43 proteins), i.e. under peroxisome-proliferating conditions when peroxisomal activity is required (**Fig. 2c, left; Tables 1, S1**). In addition, the peroxins Pex9, Pex12, and Pex15 were exclusively identified in oleate-grown cells. Pex9, a paralog of Pex5, is an additional receptor for PTS1 proteins that is only expressed in the presence of oleate (Effelsberg et al. 2016; Yifrach et al. 2016). The RING-finger peroxin Pex12 and Pex15, the membrane anchor for the Pex1/Pex6 AAA complex, are required for functional peroxisomal matrix protein import (Elgersma et al. 1997; Albertini et al. 2001) and can therefore be expected to be present in glucose-grown cells as well. However, they are of low abundance even under peroxisome-inducing conditions (**Table 1**) and were probably below the detection limit in our study. Eight peroxisomal proteins showed a higher abundance under fermentative conditions. One protein, the glutathione transferase Gto1, was only detected under these conditions. The remaining 48 peroxisomal proteins identified in our study did not show significant differences between the growth conditions.

The highest oleate-to-glucose ratios, reaching up to > 500-fold, were observed for peroxisomal matrix proteins, in particular enzymes of the fatty acid beta-oxidation and the glyoxylate cycle (**Figs. 2d, S2b, S2c**; **Table 1**). Many of these enzymes are negatively regulated by glucose-repression in glucose-grown cells (see Hiltunen et al. 2003 and references therein). Furthermore, the expression of most genes coding for enzymes of the beta-oxidation pathway and the glyoxylate cycle is induced under oleate by transcriptional activation through the Pip2-Oaf1 transcription factor, which binds to oleate-responsive elements in the promoter region of the genes (Karpichev et al. 1997; Hiltunen et al. 2003). In addition to these enzymes, levels of peroxisomal membrane and membrane-associated proteins (e.g. peroxins, transporters, as well as peroxisomal proliferation, division, inheritance, contact site or quality control factors) were ∼20 to 60- fold higher in oleate (**Figs. 2e, 2f, S2d-f; Table 1**).

Despite the drastically increased levels of peroxisomal enzymes in oleate- compared to glucose-grown cells (**Figs. 2d, S2b, S2c**; **Table 1**), changes in the levels of peroxins involved in matrix protein import are moderate (**Fig. 2f**; **Table 1**). The cytosolic receptor for PTS1 proteins, Pex5, showed a ∼6-fold higher abundance in oleate, while the import factors for PTS2 proteins remained unchanged (Pex7) or were ∼2-fold increased (Pex18 and Pex39; Pex21 was not detected). Furthermore, the components of the peroxisomal docking complex (Pex14, Pex13, Pex17) were ∼2- to 8-fold, and proteins of the Pex5 export machinery (Pex22, Pex4, Pex1, Pex6,) ∼2- to 6-fold increased in oleate-grown cells.

In addition to glutathione transferase Gto1 (detected only in glucose), a few peroxisomal proteins showed a higher abundance under fermentative conditions. These include the enzymes Tsr3, Pre2, Glr1 and Lpx2 (**Fig. 2d**), which only partially localize to peroxisomes (Yifrach et al. 2022), the very long-chain fatty acid transporter Fat1 (**Figs. 2e, S2d**), the nicotinamidase Pnc1 (**Fig. 2d**), and two proteins involved in amino acid metabolism (Str3 and Lys4; **Fig. 2d**). The latter observation is in line with the results of the GOBP term enrichment analysis showing that proteins of amino acid biosynthesis processes are enriched in glucose-grown cells (**Fig. 2b, Table S2**).

To conclude, mainly enzymes involved in fatty acid beta-oxidation and the glyoxylate cycle are drastically increased under oleate conditions, which reflects the regulation of gene expression by glucose-repression and/or oleate-induction. While our data show that the peroxisomal protein import machinery adapts to some degree to the load of newly synthesized peroxisomal enzymes at the expression level, it also appears to provide a high range of import capacity to meet the metabolic demands of cells.

### Absolute quantitative dimension of the peroxisomal proteome

Our protein copy number estimation allows us to add an absolute quantitative dimension in terms of protein copies/molecules per cell to the relative differences in protein abundance between the two metabolic conditions. Peroxisomal proteins constitute 2.8% of the total cellular proteome of cells grown on oleate, corresponding to 2.01 x 10^6^ proteins. In comparison, mitochondrial proteins are ∼11-fold higher in total abundance, despite the fact that oleate is exclusively metabolized in peroxisomes to produce energy and metabolites for cell survival and growth under these conditions. The peroxisomal proteome of glucose-grown cells amounts to 0.8% of the entire proteome, i.e. 6.67 x 10^5^ proteins. Under these conditions, the abundance of the mitochondrial proteome is ∼33- fold higher (**Fig. 3a**, see also **Fig. 1b right**).

**Fig. 3.**
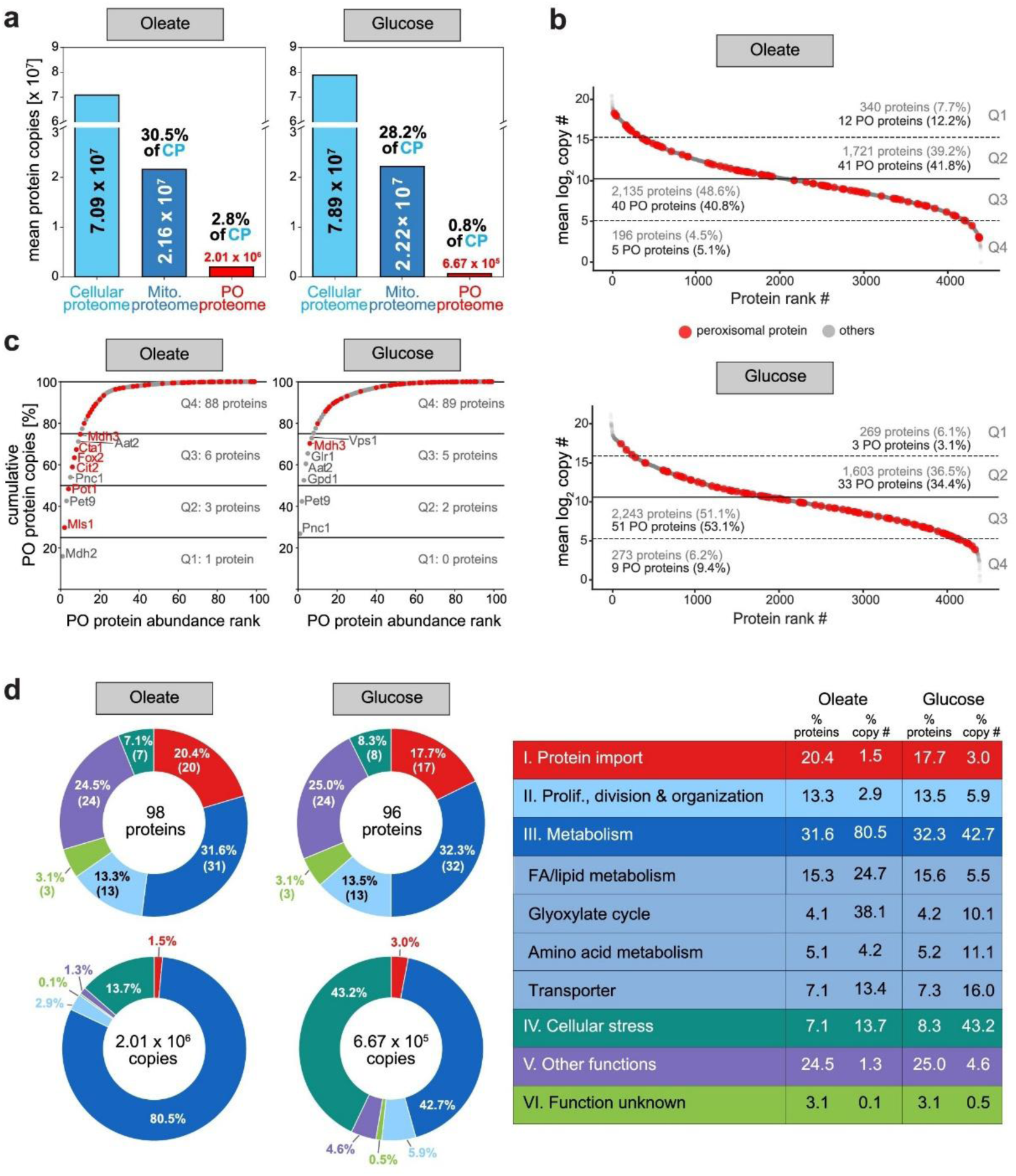
The peroxisomal proteome at absolute quantitative scale. **a.** Size of the peroxisomal proteome compared to the entire cellular proteome (CP) and the mitochondrial (Mito.) proteome under peroxisome-inducing (oleate) and fermentative (glucose) conditions. **b.** Dynamic range of the cellular and the peroxisomal proteome (marked in red) in oleate- and glucose-grown cells. The cellular proteome was divided into four quantiles (Q1 – Q4) based on mean log_2_ copy numbers with Q1 containing the proteins with highest and Q4 the proteins with the lowest copy numbers. PO, peroxisomal. **c.** Cumulative peroxisomal protein copy number plots. Proteins exclusively or predominantly localized in peroxisomes are marked in red. **d.** Fraction of proteins of different functional groups as indicated in the peroxisomal proteome of oleate- and glucose-grown cells, based on both the number of distinct proteins and their abundance (i.e., protein copy numbers). Number in brackets (donut charts, top) indicate number of proteins assigned to this functional group.

To assess the distribution of all cellular proteins depending on their copy numbers across the entire range, we divided the copy number data for oleate (range: 1 - 1.44 x 10^6^ copies per protein) and glucose (1 - 2.41 x 10^6^ copies) into four quantiles based on mean log_2_ copy numbers. The quantile comprising the proteins with the highest copy numbers (Q1) contained 340 proteins (7.7%) in oleate- and 269 proteins (6.1%) in glucose-grown cells, while 196 proteins (4.5%, oleate) and 273 proteins (6.2%, glucose) accounted for the low abundance proteome (Q4) covered in our study (**Fig. 3b, Table S1**). The remaining ∼ 88% of the proteins in both oleate- and glucose-grown cells were distributed across Q2 and Q3, with higher numbers in Q3. Peroxisomal proteins span a dynamic range of 8 – 319,537 copies in oleate- and 15 – 178,852 copies in glucose-grown cells (**Table 1**), considering both the peroxisomal core proteome (i.e., all peroxins plus further proteins exclusively or predominantly localized to peroxisomes) and multi-localized proteins. In oleate-grown cells, peroxisomal proteins are shifted to the high abundance proteome, with 53 proteins (54%) present in Q4 and Q3 compared to 36 proteins (37.5%) in glucose-grown cells (**Fig. 3b**). Ten of the 98 peroxisomal proteins identified in oleate-grown cells make up ∼75% of all peroxisomal protein copy numbers. These proteins include enzymes of the beta-oxidation pathway (Pot1, Fox2, Cta1) and the glyoxylate cycle (Mdh2, Mls1, Cit2, Mdh3), the main metabolic pathways under these growth conditions, as well as three proteins of other function (Pex9, Pnc1, Aat2), which are not exclusively localized to peroxisomes (**Fig. 3c, left**). In glucose-grown cells, the mulitlocalized nicotinamidase Pnc1 alone accounts for more than 25% of the peroxisomal protein copy numbers (**Fig. 3c, right**). Together with further five proteins (Pet9, Gpd1, Aat2, Glr1, Vps1), which are only partially localized to peroxisomes, and the peroxisomal Mdh3, it makes up ∼75% of the peroxisomal protein copy numbers under fermentative growth conditions.

**Figure 3d** shows the functional classification of the peroxisomal proteome and the corresponding number of proteins and protein copy numbers for both metabolic conditions. Proteins involved in metabolism represent the largest functional class under oleate conditions, both in number of different proteins (31.6%) and copy numbers (80.6%). Under fermentative conditions, metabolic proteins and cellular stress-related proteins represent the largest fractions of all protein copies (∼43% each), albeit overall copy numbers for stress-related proteins are similar on oleate (2.72 x 10^5^) and glucose (2.86 x 10^5^). Remarkably, although proteins required for peroxisomal protein import account for ∼20% of all peroxisomal proteins under both growth conditions (oleate, 20.4%; glucose, 17.7%), they only make up 1.5% (oleate) and 3.0% (glucose) of the peroxisomal protein copy numbers. Peroxisomal proteins with other known functions include proteins involved in heme biosynthesis (Hem14), ribosome biogenesis (Nan1), tRNA and rRNA processing (Ncs6, Iki1, Tad3, Tsr3), protein ubiquitination (Gid7, Rmd5), to name just a few. They make up ∼24% of the peroxisomal proteome in number but only 1 - 4% in absolute abundance. However, a function in peroxisome biology still needs to be established for these proteins. Peroxisomal proteins of so far unknown function contribute to ∼5% in number and are of low abundance. In the following, we will discuss the results for individual proteins of different functional classes in detail.

### Proteins involved in lipid metabolism and the glyoxylate cycle

Proteins functioning in lipid metabolism, in particular fatty acid beta-oxidation, and the glyoxylate cycle show a drastic increase in copy numbers in oleate- versus glucose-grown cells (**Figs. 3d, S3; Table 1**). Summed copy numbers are ∼13-fold higher on oleate (1.34 x 10^6^ copies per cell) than on glucose (1.04 x 10^5^ copies per cell), accounting for 67% and 16% of the entire peroxisomal proteome in these cells. These numbers illustrate the dynamics of the peroxisomal proteome in dependence of the carbon source at an absolute quantitative scale.

**Figure 4** illustrates the protein copy number data for the enzymes involved in lipid/fatty acid metabolism and the glyoxylate cycle at a single protein level. Medium chain fatty acids (MCFAs, chain length C6 – C12) presumably pass the peroxisomal membrane by diffusion (van Roermund et al. 2012). Inside peroxisomes, they are activated to acyl-CoA esters by the acyl-CoA synthetase Faa2 (Palmieri et al. 2001). The copy numbers of Faa2 (32,257 oleate/521 glucose) reflect its strong induction under oleate condition and qualifies it as one of the most abundant proteins in peroxisomes. In contrast to MCFAs, oleate and other long chain fatty acids (LCFA; chain length C13 – C21) are activated outside of peroxisomes and need the heterodimeric peroxisomal ABC transport complex Pxa1/Pxa2 for peroxisomal import (Hettema et al. 1996). Interestingly, copy numbers for Pxa2 were ∼11-fold higher in oleate-grown cells (801 vs. 75 copies in glucose), whereas the absolute abundance of Pxa1 was only 1.5-fold increased (424 versus 279 copies). While the higher copy numbers of Pxa1 and Pxa2 in oleate-grown cells reflect the need for increased import of LCFAs as substrate for peroxisomal fatty acid beta-oxidation, our data also indicates that the ratio of Pxa1 to Pxa2 is ∼1:2. Thus, the import of LCFAs in particular depends on a strong induction of Pxa2 in oleate condition but might be limited by Pxa1 levels. However, calculated copy numbers are generally estimates and further investigation is needed to confirm these differences in Pxa1/Pxa2 abundance and the functional implications.

**Fig. 4.**
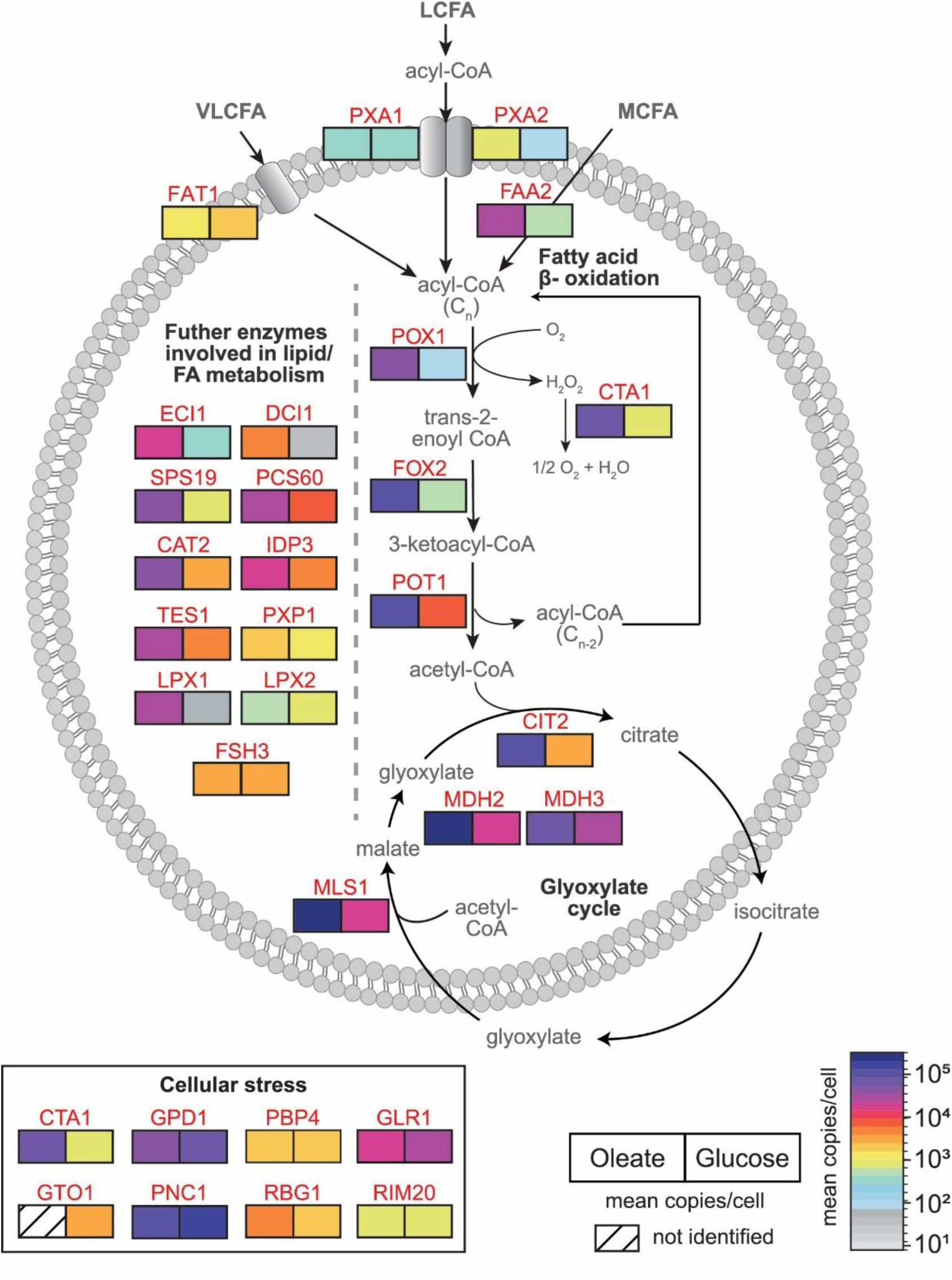
Absolute quantitative map of peroxisomal proteins associated with selected metabolic and stress-related processes. Shown are proteins of the peroxisomal fatty acid and lipid metabolism, including fatty acid beta-oxidation, and peroxisomal proteins with known or potential implication in cellular stress response and their carbon source-dependent mean copy numbers per cell and carbon source (n=4 each).

Copy numbers of FAT1, an acyl-CoA synthetase with a proposed role in the transport of very long-chain fatty acids (chain-length ≥ C22) (Zou et al. 2002), were slightly lower under oleate (1,048) than glucose (1,758). This observation is consistent with previous reports showing that the acyl-CoA synthetase activity of Fat1 is higher when cells are grown in glucose compared to oleate (Choi and Martin 1999), which could be a direct result of the differences in Fat1 protein levels.

Following import, acyl-CoA esters are degraded to acetyl-CoA and an acyl-CoA ester shortened by two carbon atoms in a series of reactions that are catalyzed by (1) the fatty-acyl coenzyme A oxidase Pox1 (Dmochowska et al. 1990; Hettema et al. 1996), (2) the multifunctional enzyme with 3-hydroxyacyl-CoA dehydrogenase and enoyl-CoA hydratase activity Fox2 (Hiltunen et al. 1992), and (3) the thiolase Pot1 (Igual et al. 1991). H_2_O_2_, generated during dehydrogenation of fatty acid acyl-CoA to trans-2-enoyl-CoA, is degraded by the catalase Cta1. All four proteins are strongly induced and part of the high abundance proteome (Q1; **Figs. 3b, S3**) of oleate-grown cells, with 51,385 copies per cell for Pox1, 89,542 for Fox2, 116,065 for Pot1 and 79,173 for Cta1. Our copy number data therefore reflect and underscore the essential function of the beta oxidation pathway, which is a hallmark of peroxisomal functions in *S. cerevisiae* cells.

Further auxiliary enzymes required for the complete degradation of unsaturated fatty acids such as oleic acid (C18:1) include Eci1 (Δ3-Δ2-enoyl-CoA isomerase), its paralog Dci1 (Δ3,5-Δ2,4-dienoyl-CoA isomerase), and Sps19 (2,4-dienoyl-CoA reductase), which were all induced under oleate. Notably, copy numbers of Dci1 (4,659 oleate/34 glucose) were considerably lower compared to Eci1 (20,883 oleate/423 glucose) independent of which carbon source was used.

Acetyl-CoA, the end product of the fatty acid beta oxidation, can be converted to acetyl- carnitine by the carnitine O-acetyltransferase Cat2 for the subsequent transfer to the cytosol and mitochondria (van Roermund et al. 1999). Cat2 is dually localized in peroxisomes and mitochondria (Elgersma et al. 1995), with ∼15-fold higher abundance in oleate (37,451 copies) versus glucose (2,574 copies) (**Fig. 4**; **Table 1**). However, under peroxisome inducing conditions, acetyl-CoA is mainly fed into the glyoxylate cycle to generate essential carbohydrates (Kunze et al. 2006). The reactions of the glyoxylate cycle are partitioned between the peroxisomal matrix and the cytosol and require the activities of citrate synthase 2 (Cit2), malate synthase 1 (Mls1), and malate dehydrogenases (Mdh3, Mdh2) in peroxisomes and aconitase 1 (Aco1) and isocitrate lyase 1 (Icl1) in the cytosol (**Fig. 4**; Minard and McAlister-Henn 1991; Chaves et al. 1997; Kunze et al. 2002; Regev-Rudzki et al. 2005; Schummer et al. 2020). The peroxisomal enzymes Cit2 (98,028 copies), Mls1 (277,581 copies), and Mdh2 (319,537 copies), and also cytosolic Icl1 (453,786) show drastically higher copy numbers in oleate- compared to glucose-grown cells (**Fig. 2c**). They are also among the proteins with the highest copy numbers in oleate-grown cells (**Fig. S3; Table 1**), emphasizing that *S. cerevisiae* cells depend on fatty acid degradation by the beta-oxidation pathway and the glyoxylate cycle under this condition.

### Cellular stress-related proteins

It is well established that peroxisomes play an important role in the response to cellular stress (Kumar et al. 2024), most notably oxidative stress (Lismont et al. 2019). Peroxisomes by definition harbor H_2_O_2_-producing oxidases and catalase to rapidly degrade the H_2_O_2_, which is toxic to cells in higher concentration. As described above, the catalase Cta1 is strongly induced in oleate-grown cells (110-fold compared to glucose) and is among the top 10 most abundant peroxisomal proteins under these conditions (79,173 copies in oleate, 714 in glucose; **Figs. 2c, 4, S3**; **Table 1**), indicative of its important function in the protection of cells from oxidative damage.

Glycerol-3-phosphate dehydrogenase 1 (Gpd1) and the nicotinamidase Pnc1 are peroxisomal enzymes whose expression is upregulated in response to different types of cellular stress (Kumar Choudhry et al. 2016). Gpd1, a key enzyme of the glycerol synthesis, is required for growth under osmotic stress (Albertyn et al. 1994), while Pnc1 is part of the NAD^+^ salvage pathway (Ghislain et al. 2002) and induced under mild heat stress or calorie restriction (Anderson et al. 2003). Our copy number data show that Gpd1 and Pnc1 are highly abundant under both growth conditions, with slightly higher levels in glucose-grown cells (Gpd1: 52856/67723 copies in oleate/glucose; Pnc1: 113,510/178852). In fact, they are the highest (Pnc1) and third highest (Gpd1) abundant peroxisomal protein in glucose (**Figs. 2c, 4, S3**; **Table 1**). Under non-stress condition, Gpd1 and Pnc1 are mainly localized to peroxisomes, with only a small fraction present in the cytosol (observed in glucose) whereas under stress, both enzymes are largely cytosolic (Anderson et al. 2003; Jung et al. 2010). Thus, peroxisomes can be considered as a “storage” of these stress-related enzymes, which can be directly released in the cytosol on demand. Peroxisomal import of both enzymes occurs via the PTS2 import route, with Pnc1, which lacks a PTS, piggy-backing on the PTS2 protein Gpd1 (Effelsberg et al. 2015; Kumar Choudhry et al. 2016).

Gto1, an omega-class glutathione transferase with glutaredoxin activity, is another enzyme associated with oxidative stress. Its expression is increased in response to oxidant conditions induced by diamide or 1-chloro-2,4-dinitrobenzene, and a role for Gto1 in the metabolism of sulfur amino acids has been proposed (Garcerá et al. 2006; Barreto et al. 2006). We identified Gto1 only in cells grown in glucose as part of the medium abundance proteome with 2,739 copies per cells (**Figs. 4, S3; Table 1**). Previous reports showed that Gto1 levels are lower in oleate- than glucose-grown cells (Barreto et al. 2006) indicating that its abundance was too low to be detected in oleate in our study. Further enzymes with potential implication in cellular stress response are the glutathione oxidoreductase Glr1, the Pbp1-binding protein Pbp4, the ribosome-interacting GTPase Rbg1, and Rim20, which were all demonstrated to be localized to peroxisomes in a recent genome-wide high-content fluorescence microscopy screen (Yifrach et al. 2022). Glr1 is required for the maintenance of the redox environment in peroxisomes (Ayer et al. 2012) and is present with 21,387 (oleate) and 33,090 (glucose) copies per cell (**Figs. 4, S3; Table 1**). In addition to peroxisomes, Glr1 is localized in the cytosol and in mitochondria (Outten and Culotta 2004). Pbp4 (4,149/2,024 copies in oleate/glucose) and Rbg1 (4,259/2,150 copies in oleate/glucose) are components of stress granules in yeast (Jain et al. 2016), suggesting a function in stress response, and Rim20 (903/972 copies in oleate/glucose; **Figs. 4, S3; Table 1**) is part of the alkaline pH response pathway (Xu and Mitchell 2001). However, the connection of Rbg1 and Rim20 to peroxisomal processes has not been shown yet.

### Proteins involved in peroxisomal matrix and membrane protein import

Import of peroxisomal matrix proteins relies on several protein complexes that form and interact with each other in a dynamic manner (see recent review by Rudowitz and Erdmann 2023). Peroxisomal matrix proteins are recognized in the cytosol by Pex5, the receptor for cargo proteins with a C-terminal peroxisomal targeting signal of type 1 (PTS1), or by Pex7, which binds cargo proteins with an N-terminal PTS2. Pex7-mediated import further requires a co-receptor (Pex18 or Pex21) for cargo binding and the recently identified biogenesis factor Pex39 to stabilize the Pex7-cargo complex prior to co-receptor binding (Chen et al. 2025).

Except for the PTS2 co-receptor Pex21, we determined the copy numbers of all known components of the cytosolic PTS1 and PTS2 cargo recognition machinery, including the alternative, oleate-inducible PTS1 receptor Pex9 (Effelsberg et al. 2016; Yifrach et al. 2016) in oleate-grown cells (**Table 1**, **Fig. 5**). In accordance with the drastically higher copy numbers of many matrix proteins in oleate versus glucose condition (see **Fig. 2d**), Pex5 has a 5.6-fold higher copy number in oleate-grown cells, but this amounts to only 274 protein copies per cell (**Table 1**, **Figs. 2f, 5, S3**). Considering that some of its cargos have 100,000 copies per cell or more (see **Figs. 4, S3,** ‘FA/lipid metabolism’ and ‘glyoxylates cycle’; **Table 1**), Pex5 receptor recycling can be seen as a resource-efficient cellular mechanism to adjust to an extremely high demand for matrix protein import as opposed to an upregulation of *PEX5* gene expression. In contrast, Pex7 has similar protein copies (∼400) under both metabolic conditions, although steady-state levels of the PTS2 protein Pot1 (imported as a dimer; Glover et al. 1994) are 16-fold higher, reaching more than 116,000 copies per cell in oleate condition (**Table 1**). Based on the difference in copy numbers of Pex5 and Pex7, it is tempting to speculate that the recycling of Pex7 is less efficient compared to Pex5 and a higher copy number of Pex7 per cell is therefore needed. It has previously been shown that recycling of Pex7 depends on Pex13 (Skowyra and Rapoport 2025), as well as the export of Pex5 and the overall process is slower than Pex5 recycling (Rodrigues et al. 2014; Hagstrom et al. 2014). Copy numbers of Pex18 (224 copies), the PTS2 co-receptor for Pot1 import, and Pex39 (118 copies) are lower than Pex7 copies, with a 1.6- to 1.8-fold increase in oleate- versus glucose-grown cells (**Table 1**). Remarkably, balanced low levels of Pex39 are critical as PTS2 protein import is not only impaired when Pex39 is lacking but also when Pex39 is overexpressed (Chen et al. 2025). Thus, the lower levels of Pex39 compared to Pex7 and Pex18 appear to not limit PTS2 protein import.

**Fig. 5.**
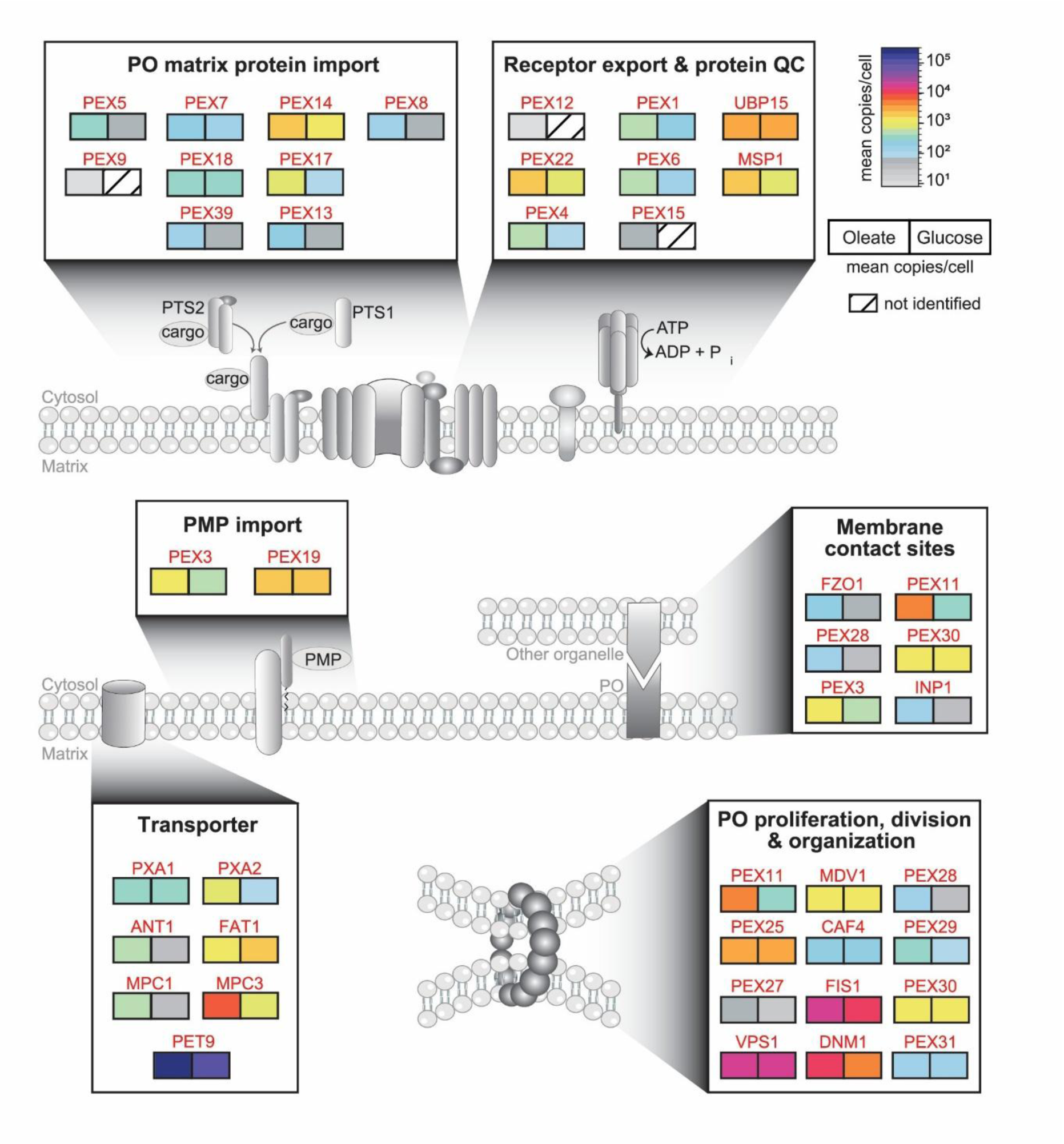
Absolute quantitative map of proteins involved in processes associated with the peroxisomal membrane. Shown are proteins of the peroxisomal (PO) matrix and membrane protein (PMP) import, receptor export and protein quality control (QC), peroxisome-associated membrane contact sites, proliferation, division and organization, and transporter/carrier proteins and their carbon source-dependent mean copy numbers per cell and carbon source (n=4 each).

At the peroxisomal membrane, receptor-cargo complexes bind to the docking complex consisting of Pex14, Pex13, Pex17, and Dyn2 (Chan et al. 2016). Absolute protein copies can be used to determine the stoichiometry of proteins in a complex. To exemplify, the Pex14/Pex17 complex that binds the cargo-loaded receptors has been shown to be assembled in a 3:1 heterotetramer (Lill et al. 2020). Our copy numbers for Pex14 and Pex17 in oleate-grown cells, 2,389 copies for Pex14 and 795 for Pex17 (**Fig. 5**; **Table 1**), agree with a 3:1 stoichiometry of this complex.

For the transfer of PTS1 and PTS2 cargo proteins into peroxisomes, Pex13 forms a conduit in the peroxisomal membrane through which the cargo is delivered by Pex5 or Pex7 and its coreceptor Pex18 or Pex21 (Gao et al. 2022; Ravindran et al. 2023; Skowyra and Rapoport 2025; Chen et al. 2025). It is estimated that ∼12 copies of Pex13 form the conduit to translocate receptor-cargo complexes into the peroxisomal lumen. Based on our data, we estimate that ∼200 Pex13 copies are present per cell under oleate and ∼55 under glucose condition. With an average of five peroxisomes present in a yeast cell grown under oleate condition, this would equal to an average number of ∼3 Pex13 conduits (with 12 copies) per peroxisome.

While it has been proposed that, after cargo release, Pex7 is retrotranslocated through the Pex13 conduit with the help of Pex39 (Skowyra and Rapoport 2025), recycling of Pex18 and Pex5 requires the ubiquitination machinery consisting of Uba1 (E1 enzyme), Pex4 (E2 enzyme, linked to the membrane via Pex22), and the RING-finger complex (Pex2/Pex10/Pex12; E3 enzyme) (Rudowitz and Erdmann 2023). Copy numbers of Pex4 and Pex22 are in the medium range and increase from 113 to 622 and 991 to 1,716 in glucose- compared to oleate-grown yeast cells (**Figs. 5, S3; Table 1**). Since one copy of Pex22 anchors one copy of Pex4 at the peroxisomal membrane (Traver et al. 2022), it is puzzling that Pex22 is ∼3-fold higher abundant than Pex4 in oleate condition. In contrast, the RING-finger proteins are of very low abundance, and only for Pex12 were we able to estimate 8 copies per cell in oleate condition. It was recently shown that the Pex2/Pex10/Pex12 complex functions as a retrotranslocation channel for Pex5 (Feng et al. 2022). Pex8 links the docking and the RING-finger complex at the luminal side of the membrane (Agne et al. 2003). It is a rather low abundant peroxin with 168 copies per cell in oleate and 60 in glucose condition (**Figs. 5, S3; Table 1**). Most recently, it has been reported that Pex8 is required for PTS1 and PTS2 cargo translocation and release in the peroxisomal matrix and that Pex8 mediates the docking of Pex5 to the RING-finger complex for receptor recycling (Ekal et al. 2025). Export of monoubiquitinated (signal for recycling) or polyubiquitinated (signal for proteasomal degradation) Pex5 (or Pex18) from peroxisomes to the cytosol requires its extraction from the peroxisomal membrane in an ATP-dependent process facilitated by the Pex1/Pex6 AAA+ complex (reviewed in Platta et al. 2024). The AAA+ complex is a heterohexamer composed of Pex1 and Pex6 in a 1:1 stoichiometry (Rüttermann et al. 2023). In line with this, we calculated equal average copy numbers of Pex1 (635) and Pex6 (573) under oleate condition. For the membrane anchor Pex15, we estimated an average copy number of 71, which is lower than the expected value for a 3:3:1 stoichiometry of the entire Pex1/Pex6/Pex15 complex. However, copy number estimates for low abundance proteins are generally less accurate, so that they can considerably deviate from the true values. Under glucose condition, Pex15 was even below the detection limit, while Pex1/Pex6 copies were ∼3-fold reduced. Peroxins required for the import of peroxisomal membrane proteins (PMPs) are Pex3 (the membrane-bound receptor for PMPs following the direct route into the peroxisomal membrane; reviewed in (Rudowitz and Erdmann 2023), and the cytosolic PMP receptor and chaperone Pex19. In line with a higher demand for PMPs under oleate condition, Pex3 levels are ∼2.5-fold higher in oleate-grown cells (1,501 vs. 586 copies in glucose). In contrast, the cytosolic PMP receptor and chaperone Pex19 has similar copy numbers under oleate (2,080 copies) and glucose (1,774 copies) condition (**Figs. 5, S3; Table 1**). To conclude, most peroxins involved in peroxisomal matrix protein import and receptor recycling are present in the low copy range (< 800 copies per cell in oleate) and induced under peroxisome-proliferating condition, except for Pex7. In comparison, copy numbers of Pex14 and Pex22 are more than 2-fold higher (> 1,700 copies per cell in oleate), suggesting additional functions as known for Pex14 of *Hansenula polymorpha* and human cells, in which Pex14 plays a role in pexophagy (van Zutphen et al. 2008; Li et al. 2025) and acts as an anchor for microtubule important for peroxisome motility (Bharti et al. 2011; Reuter et al. 2021). Furthermore, copy numbers of Pex3 and Pex19 are considerably high, with a 3-fold induction of Pex3 but not Pex19 under oleate. Pex3 is also involved in pexophagy, membrane contact sites, and inheritance (Munck et al. 2009; Motley et al. 2012; Knoblach et al. 2013; Hulmes et al. 2020), which might explain its rather high absolute abundance in yeast cells.

### Transporter/carrier proteins

Peroxisomes are metabolic organelles that need to exchange metabolites and co-factors across the membrane to function properly. In this study, we identified seven transporter/carrier proteins with known peroxisomal localization (Pxa1, Pxa2, Ant1, Fat1, Pet9, Mpc1 and Mpc3). All transporter/carrier proteins have higher copy numbers in oleate- than in glucose-grown cells, except for Fat1 (**Figs. 4, 5, S3**; **Table 1**). Besides the transporters (Pxa1, Pxa2, Fat1) with a role in fatty acid beta-oxidation (discussed above), known peroxisomal transporters have roles in nucleotide and pyruvate transport. Ant1 is an adenine nucleotide transporter that mediates the exchange of cytosolic ATP for peroxisomal AMP or ADP (van Roermund et al. 2001; Palmieri et al. 2001). Copy numbers of Ant1 are ∼17-fold increased under oleate (655 *versus* 38 copies in glucose), which reflects the high metabolic activity of peroxisomes under this growth condition. Different to peroxisomal Ant1, the ADP/ATP carrier Pet9 (alias Aac2) and the pyruvate carriers Mpc1 and Mpc3 are mitochondrial carrier proteins that have recently been shown to be partially localized in peroxisomes (van Roermund et al. 2021; Kosir et al. 2025), all having higher copy numbers in oleate-grown cells. Pet9 is present in the high abundance proteome in both oleate- and glucose-grown cells (258,248 copies on oleate vs. 103,862 on glucose), while Mpc1 and Mpc3 are of lower abundance but with ∼10-fold increase in oleate-grown cells (495 vs. 38 copies for Mpc1, 7992 vs. 783 copies for Mpc3). However, whether the higher copy numbers observed for these three transporters under glucose derepression conditions reflect a higher metabolic activity of mitochondria or peroxisomes or both remains an open question.

### Proteins mediating peroxisome proliferation, division, and organization

Growth of yeast cells under peroxisome-proliferating conditions requires the machinery for fission of preexisting peroxisomes (reviewed in Schrader et al. 2012). The process relies on the concerted actions of Pex11 and the Pex11-related peroxins Pex25 and Pex27; the tail-anchored protein and anchor for the fission machinery Fis1 and the adaptor proteins Caf4 and Mdv1, which together recruit the dynamin-like GTPase Dnm1 to the division site of the membrane; and Vps1, another dynamin-like protein that has recently been shown to function together with Pex27 in peroxisome fission (Ekal et al. 2023). Of note, Fis1, Caf4, Mdv1 and Dnm1 are shared between peroxisomes and mitochondria (Schrader et al. 2012). Copy numbers of all these proteins span a range of 50 to 20,176 copies in oleate and 20 to 17,040 copies in glucose-grown cells, with Pex27 being the lowest and Vps1 the highest abundant protein (**Figs. 5, S3; Table 1**). In addition to peroxisome fission, Vps1 is involved in many other cellular processes including endocytosis (Smaczynska-de Rooij et al. 2010), endosomal trafficking (Wilsbach and Payne 1993; Nothwehr et al. 1996; Lukehart et al. 2013), actin cytoskeleton organization (Yu and Cai 2004), and pexophagy (Mao et al. 2014), which may be the reason for its high abundance in yeast cells. Pex11, the protein required for initiating the division process, is ∼12-fold higher abundant in oleate- than in glucose-grown cells (3,987 vs. 323 copies), which is a result of *PEX11* gene induction by oleate (Gurvitz et al. 2001). Interestingly, copy numbers of Pex25 (3,187) and Pex11 are similar in oleate, whereas the family member Pex27 is 80-fold lower in absolute abundance (50 copies). Further, Pex25 copy numbers do not change between the two carbon-sources and the increase in Pex27 copies is moderate (2.5-fold) under oleate condition. Of note, Pex11 and Pex25 are the most abundant peroxins in oleate-grown cells.

Further proteins that control peroxisome number and size in *S. cerevisiae* are the Pex30 protein family members Pex28, Pex29, Pex30, Pex31, and Pex32 (for details, see review by (Deori and Nagotu 2022), which are all ER membrane proteins (Mast et al. 2016; Joshi et al. 2016; Ferreira and Carvalho 2021). Pex32 is expressed only at a very low level, which is the reason why it was not identified in our analysis. Except for Pex30, the Pex30 family proteins have generally low copy numbers (118 to 347 in oleate- and 44 to 131 in glucose grown cells), showing slightly decreased (Pex31) or ∼3-fold higher abundance (Pex28, Pex29) in oleate-grown cells (**Figs. 5, S3; Table 1**). Pex30 copy numbers are in the medium range with slightly higher numbers in oleate (1,567) versus glucose (1,339). Taken together, the protein copy numbers for the Pex30 family proteins are rather unaffected by the carbon source, with PEX30 being the most abundant family member. In addition to their function in regulating peroxisome number and size, Pex30 family members are also engaged in different membrane contact sites, as discussed in the next section.

### Peroxisome-associated membrane contact site proteins

Peroxisomes form membrane contact sites (MCS) with various other organelles to coordinate metabolic functions, facilitate lipid and ion exchange, and maintain cellular homeostasis. MCS are regions of close appositions between membranes of the partner organelles and are mediated by protein-protein or protein-lipid interactions that physically tether the organelles (Scorrano et al. 2019). Our dataset provides copy numbers for proteins that are part of peroxisome-associated MCS (**Figs. 5, S3; Tables 1, S1**). Peroxisome-mitochondria (“PerMit”) contacts are mediated by Pex11 and Mdm34, a mitochondrial component of the ERMES complex (Mattiazzi Ušaj et al. 2015). The oleate-inducible Pex11 is present with ∼4,000 copies in oleate (323 in glucose), while Mdm34 has only 27 (oleate) and 36 (glucose) copies per cell (**Table S1**). The very low abundance of Mdm34 indicates that a Pex11-Mdm34 tether is a rather rare event under the applied conditions. Further proteins involved in PerMit contacts are Fzo1 in the mitochondrial outer membrane and the PMP Pex34 (Shai et al. 2018), which was not identified here. Copy numbers of the mitochondrial mitofusin Fzo1 considerably increased from 57 in glucose to 210 in oleate-grown cells. In line with this finding, Fzo1 was shown to form PerMit contacts via a homotypic interaction, which increases with higher fatty acid desaturation (Alsayyah et al. 2024). Under these conditions, Fzo1-PerMit contacts promote the transfer of citrate from peroxisomes to mitochondria, where it is metabolized in the tricarboxylic acid cycle to promote mitochondrial activity. Since oleic acid is an unsaturated fatty acid, we propose that Fzo1-mediated PerMit contacts are likely also increased in oleate-grown cells. This notion is underscored by the strong overrepresentation of proteins of the TCA cycle and the mitochondrial electron transfer chain under peroxisome-inducing versus fermentative growth condition (**Fig. 2b**).

As mentioned above, all five members of the Pex30 protein family have been reported to be present at MCS with various organelles (David et al. 2013; Ferreira and Carvalho 2021; Hugenroth et al. 2025) and localize in the ER membrane (Mast et al. 2016; Joshi et al. 2016; Ferreira and Carvalho 2021). Together with Pex28 and Pex32, Pex30 forms a complex at ER-peroxisome contact sites (EPCONs) (Ferreira and Carvalho 2021). If or how they form a tether between ER and peroxisomal membrane is not known since a tethering partner in the peroxisomal membrane has not been identified. However, cells lacking the *PEX30* gene are defective in ER-peroxisome contact site formation underscoring the importance of Pex30 for EPCON integrity (Ferreira et al. 2025). The higher absolute abundance of Pex30 might also be explained by its further involvement in MCS at nucleus-vacuole junctions (NVJs), via binding to Pex29 (Ferreira and Carvalho 2021), and its accumulation at ER-lipid droplet contacts (Ferreira and Carvalho 2021). While we did not identify Pex32, our data cover the low abundance EPCON protein Pex28. Copy numbers of Pex28 were 3-fold higher in oleate-grown cells (142 vs. 44 in glucose) (**Figs. 5, S3; Table 1**), pointing to a higher number of Pex28/Pex30/Pex32-dependent EPCONs under these conditions.

EPCONs are also formed by Pex3, which is present in both the ER and the peroxisomal membrane and bridged by the peroxisomal protein Inp1 (Knoblach et al. 2013). The Pex3- Inp1 protein pair also tethers peroxisomes to the plasma membrane by binding of Inp1 to Pex3 in the peroxisomal and phosphatidylinositol 4,5-bisphosphate in the plasma membrane (Hulmes et al. 2020). Both, tethering peroxisomes to the ER and the plasma membrane, is required for peroxisome retention in the mother cell during cell division as a mechanism to control the distribution of peroxisomes between mother and daughter cells). Pex3 is a rather abundant peroxin with 1,501 copies per cell (∼300 per peroxisome) in oleate, which likely reflects its multiple roles in peroxisomal membrane protein import, pexophagy and MCS. In contrast, Inp1 is a low abundance protein with 122 copies per cell in oleate and 39 in glucose (**Figs. 5, S3; Table 1**).

Taken together, our data show that MCS proteins with several functions associated with peroxisomes have copy numbers in the medium to high range (Pex11: 3,987; Pex30: 1,567; Pex3: 1,501 – all numbers for oleate), while the MCS proteins Fzo1, Pex28, Inp1 are present at low copy numbers (210 to 122 copies in oleate) or below the detection limit (Pex32) in this work. Thus, tethering events are likely dictated by these low copy factors and reflect rather specific, regulated events depending on the metabolic needs of a cell.

## Concluding Remarks

We here report absolute quantitative data in form of copy numbers for peroxisomal proteins that are expressed in cells that were grown under peroxisome-inducing in comparison to fermentative conditions. The data enable insight into a new, quantitative dimension of peroxisome biology, e.g. the dynamic range, the regulation and dynamics of the peroxisomal proteome, the capacity of the peroxisomal matrix and membrane protein import machinery, and the average number of distinct protein complexes per peroxisome. Assuming a number of five peroxisomes per oleate- and two per glucose-grown cell, a peroxisome contains ∼ 200,000 matrix protein copies under oleate compared to ∼ 46,000 copies under glucose condition, i.e. less than a quarter (only counting matrix proteins of the peroxisomal core proteome). In comparison, all peroxins (including cytosolic import factors) account for 2.12 x 10^4^ and 1.13 x 10^4^ copies in oleate and glucose condition, respectively. 160 copies of Pex14/Pex17 complexes per peroxisome on oleate and 50 on glucose provide docking sites for the PTS import receptors, and all matrix proteins are imported into peroxisomes through ∼3 (oleate) or ∼2 (glucose) Pex13 conduits. It is important to note, though, that the reported protein copy numbers are estimates, which in particular applies to low abundance proteins (< 200 copies per cell). We further envision that our data will guide the characterization of the ultrastructural arrangements of proteins in three-dimensional reconstructions of peroxisomes obtained by electron microscopy tomography. In conclusion, our absolute quantification of the peroxisomal proteome in *S. cerevisiae* will foster peroxisomes research and serves as a point of reference for the peroxisome community.

## Supporting information

Protein copy numbers per cell for S. cerevisiae grown in oleate- or glucose

Results of GO term enrichment analyses

## Author contributions

BW conceptualized and supervised the study and secured funding; RM performed the experiments; HD, SO, and JB performed data analyses and prepared figures; SO wrote the original draft of the manuscript; BW reviewed and edited the manuscript. Work included in this study has been performed in partial fulfilment of the doctoral thesis of HD and RM at the University of Würzburg.

