## Supplementary Figures S1 - S3 for "The yeast peroxisomal proteome at absolute quantitative scale"

**Supplementary information contents:**

Supplementary Fig. S1
Supplementary Fig. S2
Supplementary Fig. S3
Legend to Supplementary Table S1
Legend to Supplementary Table S2

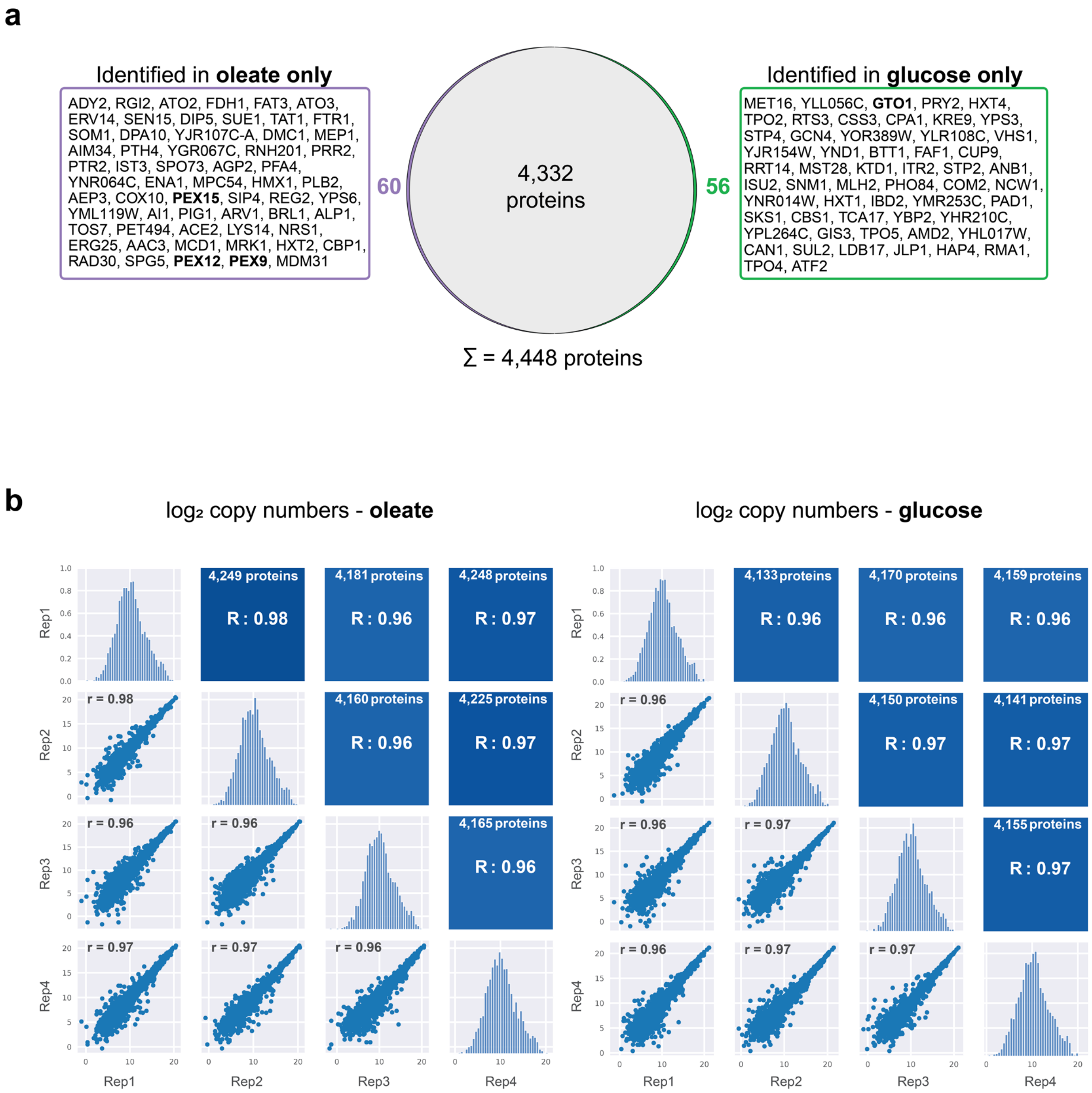


**Supplementary Fig. S1 | Evaluation of cellular protein copy number estimation**

Related to figure 1. **a.** Overlap of proteins identified in oleate- and glucose-grown cells. Proteins identified only under one of the conditions are listed; peroxisomal proteins are marked in bold. **b.** Multiscatterplot showing the reproducibility of protein copy numbers ($log_2$-transformed values) determined for individual replicates (Rep) of oleate- (left) and glucose-grown (right) cells (n=4 replicates each).

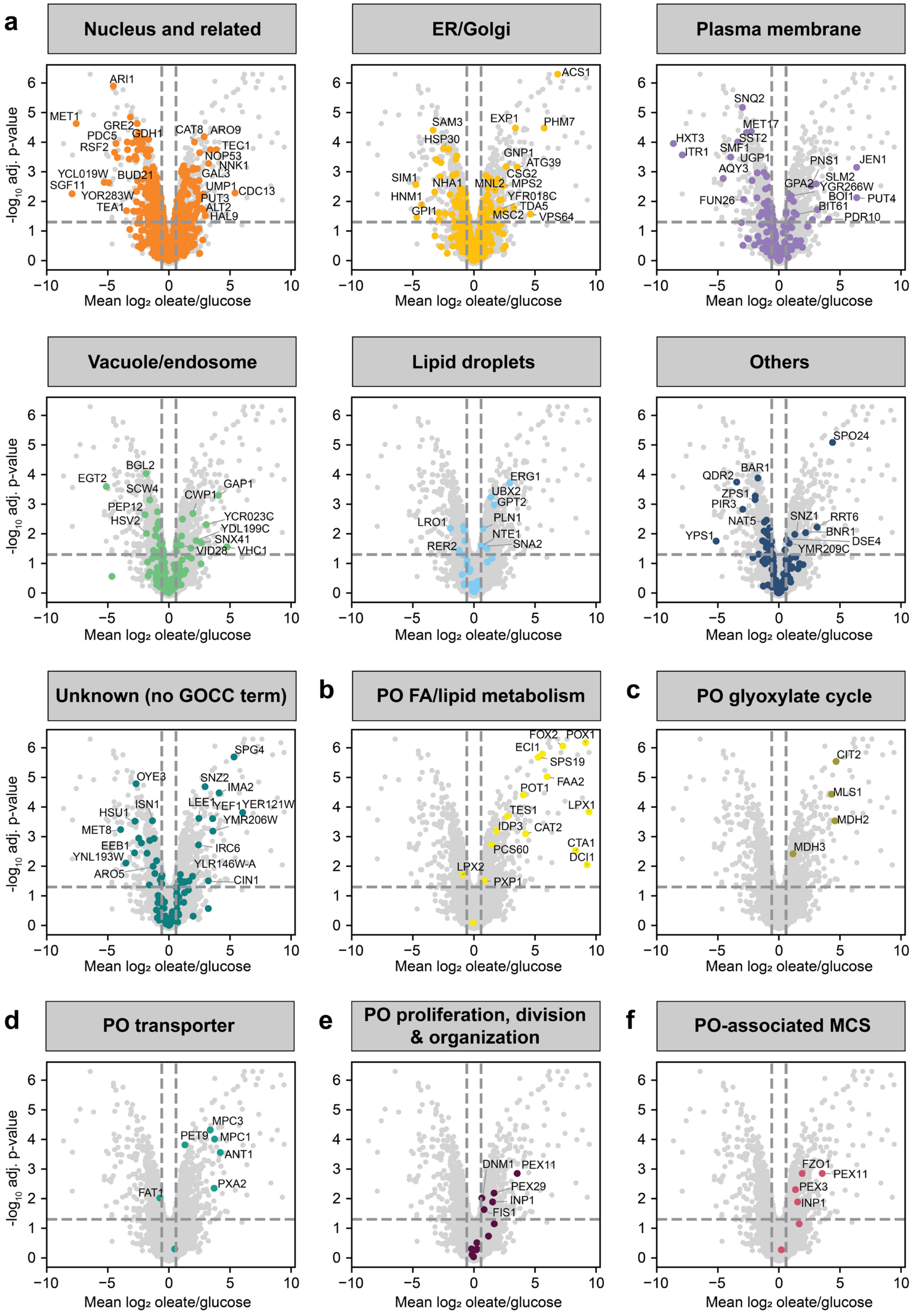
a
Nucleus and related
ARI1
MET1
GRE2
PDC5
GDH1
RSF2
CAT8
ARO9
TEC1
NOP53
NNK1
YCL019W
BUD21
GAL3
SGF11
UMP1
CDC13
YOR283W
PUT3
TEA1
ALT2
HAL9
ER/Golgi
ACS1
SAM3
EXP1
PHM7
HSP30
GNP1
ATG39
SIM1
NHA1
MNL2
CSG2
MPS2
HNM1
YFR018C
TDA5
GPI1
MSC2
VPS64
Plasma membrane
SNQ2
MET17
HXT3
SST2
ITR1
SMF1
UGP1
AQY3
PNS1
JEN1
SLM2
GPA2
YGR266W
FUN26
BOI1
PUT4
BIT61
PDR10
Vacuole/endosome
BGL2
EGT2
SCW4
GAP1
CWP1
PEP12
HSV2
YCR023C
YDL199C
SNX41
VID28
VHC1
Lipid droplets
ERG1
UBX2
GPT2
LRO1
PLN1
NTE1
RER2
SNA2
Others
SPO24
BAR1
QDR2
ZPS1
PIR3
NAT5
SNZ1
RRT6
YPS1
BNR1
DSE4
YMR209C
Unknown (no GOCC term)
SPG4
OYE3
SNZ2
IMA2
ISN1
LEE1
YEF1
YER121W
HSU1
YMR206W
MET8
EEB1
IRC6
YNL193W
YLR146W-A
ARO5
CIN1
b
PO FA/lipid metabolism
FOX2
POX1
ECI1
SPS19
FAA2
POT1
LPX1
TES1
IDP3
CAT2
PCS60
CTA1
DCI1
LPX2
PXP1
c
PO glyoxylate cycle
CIT2
MLS1
MDH2
MDH3
d
PO transporter
MPC3
PET9
MPC1
ANT1
PXA2
FAT1
e
PO proliferation, division & organization
DNM1
PEX11
PEX29
INP1
FIS1
f
PO-associated MCS
FZO1
PEX11
PEX3
INP1
-log10 adj. p-value
Mean log2 oleate/glucose
−10
−5
0
5
10
0
1
2
3
4
5
6

**Supplementary Fig. S2 | Carbon source-dependent differences in the abundance of proteins of different subcellular localization and associated with different peroxiso-mal processes**

Related to figure 2. **a.** Same plot as shown in Figure **2a** highlighting proteins of the indicated subcellular localization (**a**) and involved in different peroxisomal processes (**b – f**) as indicated. PO, peroxisomal; FA, fatty acid; MCS, membrane contact sites.

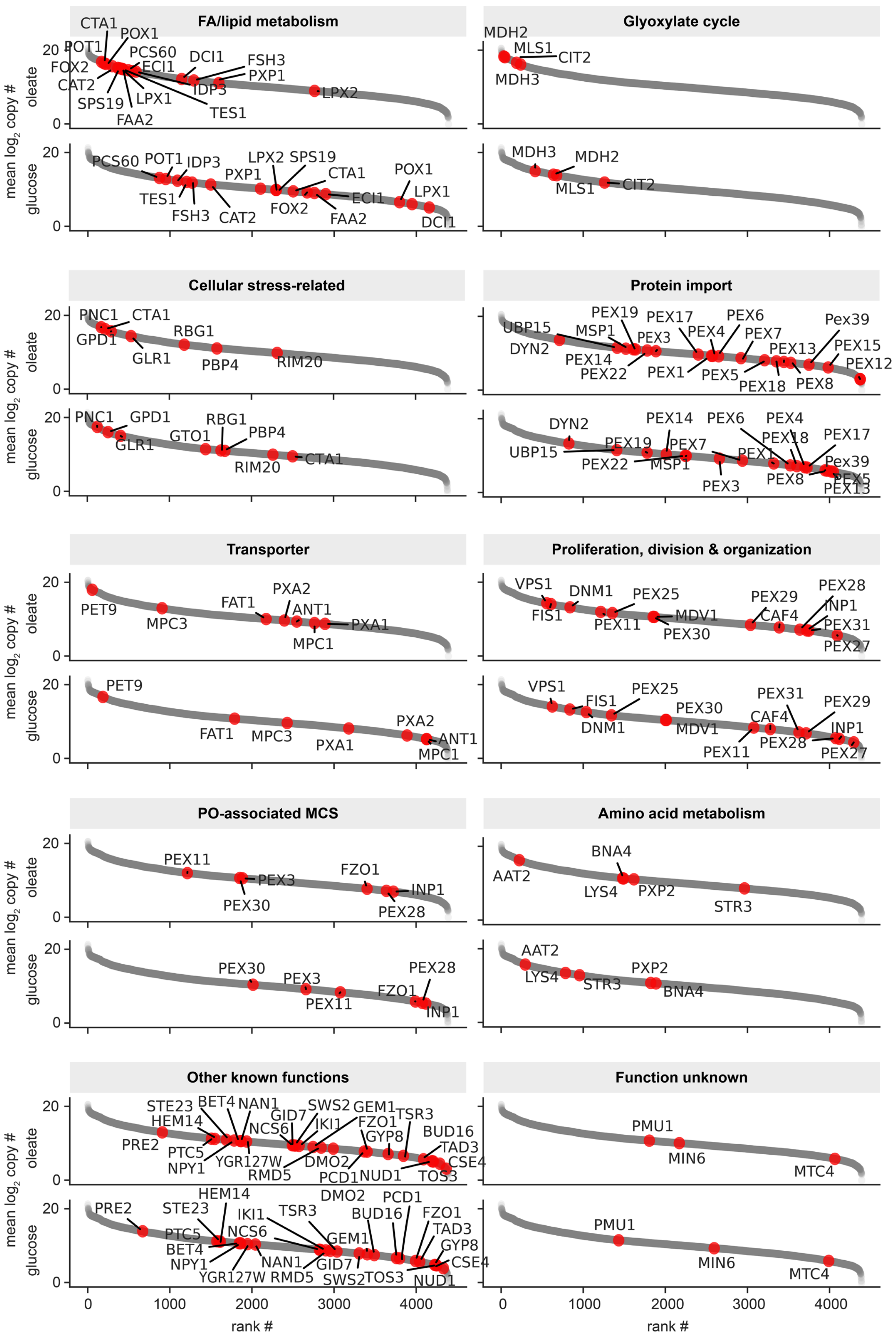

FA/lipid metabolism
Glyoxylate cycle
Cellular stress-related
Protein import
Transporter
Proliferation, division & organization
PO-associated MCS
Amino acid metabolism
Other known functions
Function unknown
mean log2 copy #
oleate
glucose
rank #
CTA1
POX1
POT1
PCS60
DCI1
FSH3
FOX2
ECI1
PXP1
CAT2
IDP3
LPX2
SPS19
LPX1
TES1
FAA2
MDH2
MLS1
CIT2
MDH3
PNC1
GPD1
GLR1
RBG1
PBP4
RIM20
GTO1
UBP15
DYN2
MSP1
PEX19
PEX17
PEX6
PEX4
PEX7
PEX3
PEX13
PEX15
PEX12
PEX14
PEX22
PEX1
PEX5
PEX18
PEX8
Pex39
PET9
MPC3
FAT1
PXA2
ANT1
PXA1
MPC1
VPS1
DNM1
PEX25
PEX29
PEX28
FIS1
PEX11
MDV1
PEX30
CAF4
INP1
PEX31
PEX27
FZO1
AAT2
BNA4
LYS4
PXP2
STR3
STE23
BET4
NAN1
GID7
SWS2
GEM1
TSR3
HEM14
NCS6
IKI1
FZO1
GYP8
BUD16
PRE2
PTC5
TAD3
NPY1
YGR127W
DMO2
CSE4
RMD5
PCD1
NUD1
TOS3
PMU1
MIN6
MTC4

**Supplementary Fig. S3 | Comparison of the absolute abundance of peroxisomal proteins in oleate- versus glucose-grown cells**

Related to figure 3. Shown are the same rank plots as in **Fig. 3b** highlighting proteins associated with the indicated peroxisomal functions. For information about the exact protein copy numbers, see **Table 1**. FA, fatty acid; MCS, membrane contact sites; PO, peroxisomal.

### Supplementary Table Legends

**Supplementary Table S1 | Protein copy numbers per cell for *S. cerevisiae* grown in oleate- or glucose** (xlsx file)

Protein copy numbers per cell were estimated based on MS1 intensities determined by MaxQuant from label-free LC-MS analyses of whole cell lysates from oleate- and glucose-grown cells ($n = 4$ each).

**Supplementary Table S2 | Results of GO term enrichment analyses** (xlsx file)

The analysis was performed for the domains "Biological Process" and "Cellular Component" for proteins with significantly higher abundance in oleate- or glucose grown cells (adjusted p-value $< 0.05$; oleate/glucose ratio $\geq 1.5$ or $\leq 0.6667$).
